# Convergent molecular changes in populations of two pipefish species in the Baltic Sea

**DOI:** 10.64898/2026.09.23.753463

**Authors:** Jule Drewalowski, Eli Taub, Oiana Trillo, Marharyta Abashkina, Gunilla Rosenqvist, Sergei Kliver, Iva Kovačić, Sarah S.T. Mak, Jessica Bogaards, Charlotta Kvarnemo, Bent Petersen, Joseph Nesme, Benoit Gouillieux, Peter Rask Møller, Rasmus Stenbak Larsen, M. Thomas P. Gilbert, Kenyon B. Mobley, Josefin Stiller

**Affiliations:** Department of Biology, University of Copenhagen, Denmark; Department of Biology, University of British Columbia, Okanagan Campus, Kelowna, BC, V1V 1V7 Canada; Department of Earth Sciences, Natural Resources and Sustainable Development, Uppsala University, Visby, Sweden; Center for Evolutionary Hologenomics, The Globe Institute, University of Copenhagen, Denmark; Department of Biological and Environmental Sciences, University of Gothenburg, Sweden; Linnaeus Center for Marine Evolutionary Biology, University of Gothenburg, Sweden; Centre of Excellence for Omics-Driven Computational Biodiscovery (COMBio), Faculty of Applied Sciences, AIMST University, Kedah, Malaysia; Department of Biotechnology and Biomedicine, Technical University of Denmark, Søltofts Plads 221, DK-2800 Kgs Lyngby, Denmark; University of Bordeaux, CNRS, Bordeaux INP, EPOC, UMR 5805, F-33600 Pessac, France; Natural History Museum Denmark, University of Copenhagen, Denmark; University Museum, NTNU, Trondheim, Norway; Norwegian College of Fishery Science, UiT The Arctic University of Norway, Tromsø, Norway

**Keywords:** Molecular convergence, Population genomics, Population structure, Baltic Sea, Syngnathidae

## Abstract

While studies of molecular convergence typically focus on species that independently evolved similar phenotypes, similar questions can be asked among populations of different species adapting to the same environment. Here, we study the Baltic Sea environmental gradient and two co-distributed, ecologically similar, but distantly related pipefish species, the broadnosed pipefish (*Syngnathus typhle*) and the straightnose pipefish (*Nerophis ophidion*), to test whether independent colonization of the Baltic Sea led to similar population structure and caused convergence at different genomic levels. Using a new diploid genome for *S. typhle* and whole-genome resequencing data (N=99 for *S. typhle*; N=81 for *N. ophidion*) across seven locations from France to Finland, we identified stronger population structure in *S. typhle* than in *N. ophidion*, while both species showed differentiation between North Atlantic, Danish Straits, and Baltic Sea populations. Genetic diversity was lower in Baltic populations, particularly in *S. typhle*. Cross-species comparisons of Baltic populations revealed contrasting chromosome-level patterns of genomic differentiation, with differentiation spread across the genome in *S. typhle* but concentrated in specific chromosomal regions in *N. ophidion*. Nonetheless, we identified multiple orthologous genes and SNPs showing convergent differentiation in the Baltic populations of both species, including genes related to metabolism, immunity, and regulatory functions. Overall, we identified molecular convergence at the gene and nucleotide level between Baltic populations of distantly related pipefish species, while convergence was limited at broader genomic scales. These findings suggest that similar environmental pressures can repeatedly target specific genetic elements even when genomic backgrounds and population structures differ.

## Introduction

A long-standing question in evolutionary biology is to which extent similar environmental conditions lead to similar adaptive changes in independent evolutionary lineages. Numerous examples of convergent and parallel evolution, such as the independent evolution of powered flight in insects, birds and bats, carnivory in pitcher plants, and echolocation in bats and toothed whales, demonstrate that shared selective pressures can lead to similar phenotypic outcomes (Losos et al. 1998; McGhee 2011). However, the extent to which phenotypic convergence is mirrored by convergence at the molecular level is less resolved.

Molecular convergence can occur at multiple hierarchical levels, ranging from individual nucleotide substitutions to repeated involvement of the same genes, regulatory networks, or functional pathways (Allard and Kumar 2026). Empirical studies have found evidence both for limited and extensive molecular convergence. In some systems, similar phenotypes evolve through multiple genetic routes (Bradley et al. 2009; Stern and Orgogozo 2009; Corbett-Detig et al. 2020), whereas in others, convergent evolution repeatedly targets the same sites, genes, regulatory networks, or functional pathways (Yokoyama and Yokoyama 1990; Zakon et al. 2006; Li et al. 2010; Liu et al. 2010; Parker et al. 2013; Chikina et al. 2016; Lyu et al. 2018; Merényi et al. 2020). Consequently, the extent of molecular convergence is expected to depend on the genetic architecture of the trait and the evolutionary divergence among lineages being compared.

Most studies of molecular convergence have focused on different species that independently evolved similar traits. Yet, similar questions arise at the population level when populations from different species adapt to similar environments (Brown et al. 2019; Greenway et al. 2024; Teweldebirhan et al. 2026). In populations, convergence depends not only on similar selective pressures imposed by the environment, but also on the amount and distribution of standing genetic variation available within each species. Because natural selection can only act on variants that are already present or newly arise within populations, species may differ in the range of genetic solutions available for adaptation (Barrett and Schluter 2008). Factors affecting population structure, such as demographic history and gene flow, shape this pool of standing variation, but it remains unclear to which extent this impacts the likelihood of convergent evolution.

The Baltic Sea provides an exceptional natural system for investigating population-level convergence. Since its formation ca. 8,000 years ago (Björck 1995), numerous marine species have independently entered this brackish inland sea from the North Atlantic, repeatedly encountering a steep environmental gradient of salinity, temperature, oxygen and light (Ojaveer et al. 2010; Bathmann et al. 2020; Geburzi et al. 2022). Salinity, for example, is at marine levels in the Skagerrak and drops quickly going east, to around 15 PSU in the Danish Straits and down to 2 PSU in the northern Baltic (Kniebusch et al. 2019; Lehmann et al. 2022). These environmental conditions pose physiological challenges impacting osmoregulation, metabolism, development, immunity and reproduction (Morgan and Iwama 1991; Haddy and Pankhurst 2000; Hill et al. 2019; Goehlich et al. 2021; Torres-Rodríguez et al. 2025) and have driven divergence in a wide range of taxa (Johannesson and André 2006; Momigliano et al. 2017; Fietz et al. 2018; Leder et al. 2021; Geburzi et al. 2022). Importantly, colonization of the Baltic Sea represents a replicated directional transition from an ancestral marine to a novel brackish environment. The repeated exposure of independent lineages to the same environmental gradient provides a natural experiment for investigating the extent to which similar selective pressures lead to convergent genomic responses.

Two species that are able to traverse the Baltic Sea environmental barrier are the broadnosed pipefish (*Syngnathus typhle* Linnaeus, 1758) and the straightnose pipefish (*Nerophis ophidion* Linnaeus, 1758), two syngnathids that diverged around 58 million years ago (Stiller et al. 2022). Both species occur in largely overlapping ranges in seagrass habitats across the Northeastern Atlantic, Mediterranean, Black Sea and Baltic Sea (Dawson 1986). Their distribution ranges in the low-salinity Baltic Sea of southern Finland indicate that they have experienced similar environmental contrasts between marine and brackish habitats, making them a suitable system to investigate molecular convergence.

Despite these shared environmental challenges, the species differ in characteristics that may influence adaptation. Both lack a dispersive larval phase due to male brooding (Whittington and Friesen 2020) and adults are poor swimmers, traits that are expected to reduce dispersal and increase population structuring (Mobley et al. 2011; Weber et al. 2015). In agreement with this expectation, *S. typhle* has been shown to have strong population structure in other parts of its range (Wilson and Eigenmann Veraguth 2010; Knutsen et al. 2022), with population showing signs of local adaptation to different salinity levels (Goehlich et al. 2021). On the other hand, *N. ophidion* may be able to disperse more than *S. typhle* because it wraps its long body around seagrass blades and macroalgae and can raft on detached parts (J. Stiller, pers. obs.), although published evidence appears to be limited. However, in contrast to *Syngnathus acus* and *Syngnathus rostellatus* that are commonly found offshore (Carl and Møller 2026), no offshore records of *N. ophidion* or *S. typhle* exists in the region. The two species differ in their genome size by five-fold (Roth et al. 2020; Ramesh et al. 2024) and differ substantially in chromosome numbers (*S. typhle*: 2n=44, *N. ophidion*: 2n=58) (Vitturi et al. 1998), suggesting distinct genomic backgrounds.

Together, these contrasts provide an opportunity to test not only whether adaptation to a shared environment results in molecular convergence, but also whether connectivity and genomic architecture influence the genomic scale at which convergence occurs. If connectivity and standing genetic variation strongly influence the evolutionary response to the Baltic Sea environment, we expect adaptive differentiation to display distinct genome-wide patterns in each species. However, if the environmental challenges in the Baltic Sea favor specific adaptive solutions, selection may nonetheless drive convergence at finer genomic scales by repeatedly targeting the same genes or even the same nucleotide variants across species.

Here, we generated a new chromosome-level genome for *S. typhle* and resequenced whole genomes for a total of 180 individuals from both species from the Northeast Atlantic into the Baltic Sea. We use this dataset to answer four questions: (i) How are populations structured within each species across the Atlantic-Baltic transition? (ii) How do the amount and distribution of standing genetic diversity available for adaptation differ? (iii) Does adaptation proceed through similar genomic architectures in the two species? and (iv) At which hierarchical genomic levels, including chromosomes, gene pathways, genes and nucleotides, do Baltic Sea populations show evidence of convergent evolution?

## Results

### Diploid genome assembly for *Syngnathus typhle* and whole-genome data for 180 individuals from seven locations for both species

We generated a chromosome-level genome for *S. typhle*, consisting of two haplotypes of 368.3 Mb and 356.3 Mb (GCA_048301445.1 and GCA_048301605.1). PacBio HiFi and Hi-C sequencing data with a coverage of 60x was used to assemble the 22 chromosomes, matching expectations from another available genome for the species (GCF_033458585.1, (Ramesh et al. 2024)) and karyotypic characterization (Vitturi et al. 1998). The genome is of high quality with N50 of 15.7 Mb and 15.3 Mb for the two haplotypes and single-copy BUSCO completeness of 96.2% and 96.0% using actinopterygii_odb10. Gene annotation for haplotype 1 (GCA_048301445.1-GB_2025_08_16) using the NCBI EGAPx pipeline identified 23,220 genes, out of which 19,756 are protein-coding.

Additionally, we produced whole-genome resequencing data for individuals of *S. typhle* (N=99) and *N. ophidion* (N=81) from seven locations along their range from the Atlantic (Arcachon, FR) to the northern Baltic (Tvärminne, FI) (Figure 1, Table S1, Table S2). Raw reads were mapped to the newly generated genome (GCA_048301605.1) from a sample from Køge (DK) and an existing chromosome-level assembly for *N. ophidion* (GCF_033978795.1, (Ramesh et al. 2024)) from an individual from Kristineberg (SE). On average, 98.4% of reads mapped (range 67.1% - 99.6%) and sequence coverage was 9x (range 1x - 35x, Table S2). Single-nucleotide polymorphisms (SNPs) were identified using genotype likelihoods, resulting in a total of 1,488,252 SNPs for *S. typhle* and 19,565,453 SNPs for *N. ophidion* (Table S3). Removing SNPs in linkage disequilibrium (LD) left 154,292 and 1,504,488 SNPs, respectively. Out of all SNPs, 264,222 in *S. typhle* and 637,432 in *N. ophidion* were located in exons. Because the total number of exonic sites are not strongly different between the two species (*S. typhle*: 36.8 Mb, *N. ophidion*, 40.7 Mb), *N. ophidion* appears to have a higher density of SNPs in the coding regions than *S. typhle*. The stark differences in SNP numbers may be explained by the roughly five times larger genome of *N. ophidion* (1,845.7 Mb) compared to *S. typhle* (356.3 Mb), while differences in mutation rate and demographic history could also contribute to differences in SNP accumulation.

**Figure 1:**
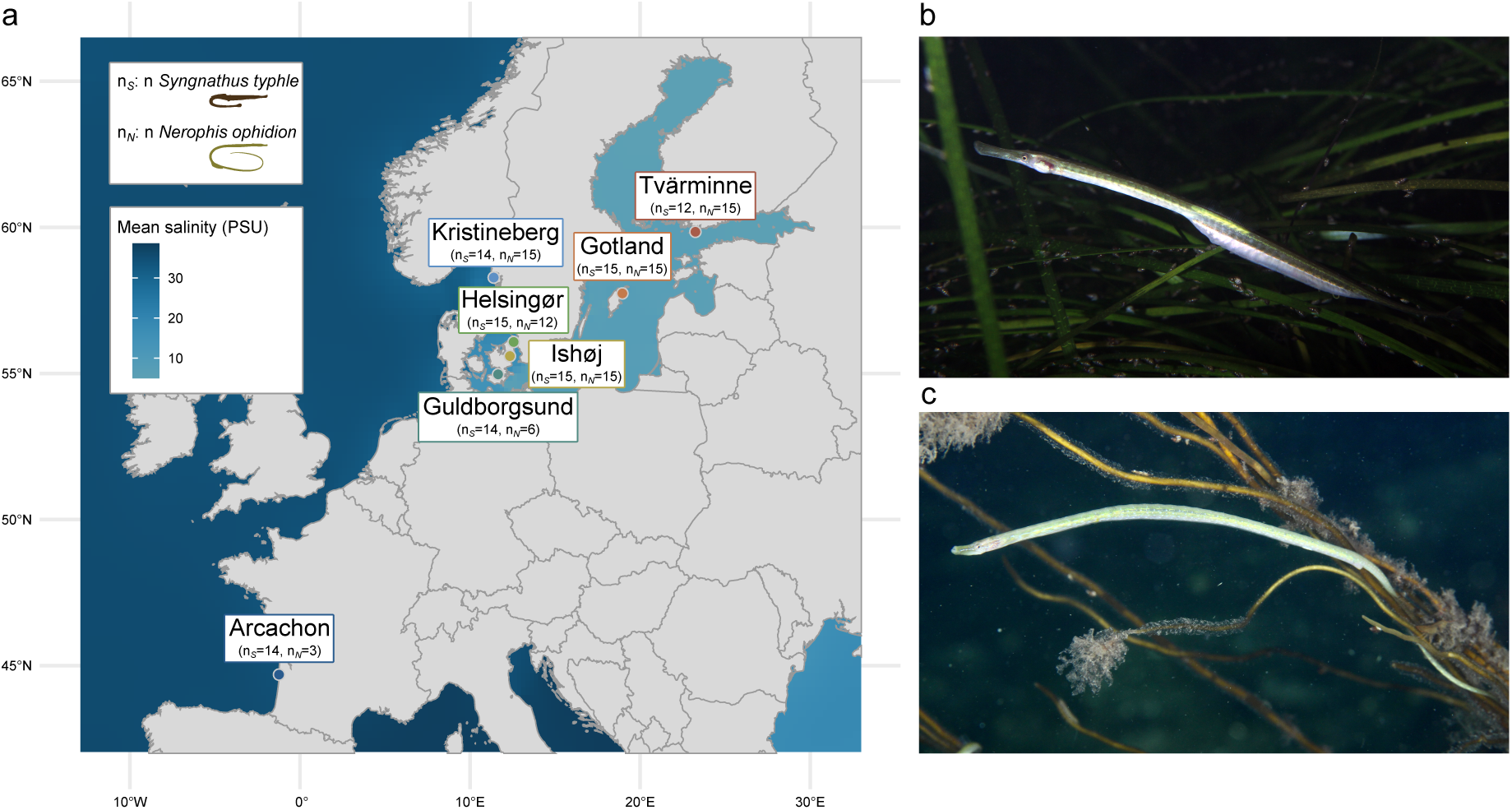
Sampling overview for two pipefish species. (a) Map of the northeast Atlantic and Baltic Sea with the water colored by mean salinity, as an example of the ecological gradient, ranging from marine salinity in the Atlantic (ca. 35 PSU) to steep salinity decrease in the Danish Straits (ca. 15 PSU) to low salinity in the Baltic (ca. 7 PSU). The seven sampling locations are marked on the map. For each location the number of sequenced individuals is given as n*_S_* for *Syngnathus typhle* and n*_N_* for *Nerophis ophidion*. Silhouette for *S. typhle* was obtained from PhyloPic, created by Alexandra Hahn; silhouette for *N. ophidion* was created for this study. Salinity data was taken from the World Ocean Atlas 2023 (Reagan et al. 2024). (b) The broadnosed pipefish (*S. typhle*) in its natural habitat. Photo provided by Peter Rask Møller. (c) The straightnose pipefish (*N. ophidion*) in its natural habitat. Photo provided by Peter Rask Møller.

### Stronger population differentiation in *Syngnathus typhle* than in *Nerophis ophidion*

Population structure differed markedly between the two species, with *S. typhle* forming three well-defined genetic groups corresponding to major geographic regions, while *N. ophidion* showed a largely continuous pattern of variation across the sampled range (Figure 2). For *S. typhle*, the PCA split samples into three clusters, namely the Atlantic (Arcachon), the Danish Straits that includes individuals collected from the Danish Straits (Guldborgsund, Helsingør, Ishøj) and Skagerrak (Kristineberg) (referred to here as the Danish straits for simplicity), and lastly the eastern Baltic Sea (Gotland, Tvärminne), subsequently referred to as Baltic Sea (Figure 2a). PC1 explained 7.72% of total variation and PC2 5.66%. PC3 clearly separated the Baltic Sea populations Gotland and Tvärminne (Figure S3a). Admixture analysis indicated that the samples were best described by five different ancestry proportions, corresponding to the three distinct clusters in the PCA, but additionally splitting Kristineberg from the remaining Danish Strait populations and splitting the Baltic Sea populations each into their own cluster (Figure 2b). Very little admixture was inferred between these five groups. Strong differentiation between the same three main clusters was also supported by the levels of genetic differentiation, as measured in F_ST_ (Figure 2c). Differentiation was largest between the Atlantic and Baltic Sea (F_ST_=0.20-0.21), relatively large between theAtlantic and the Danish Straits (F_ST_=0.09), and the Baltic Sea populations were moderately differentiated to the Danish Straits (F_ST_=0.10-0.12) and between each other (F_ST_=0.08).

**Figure 2:**
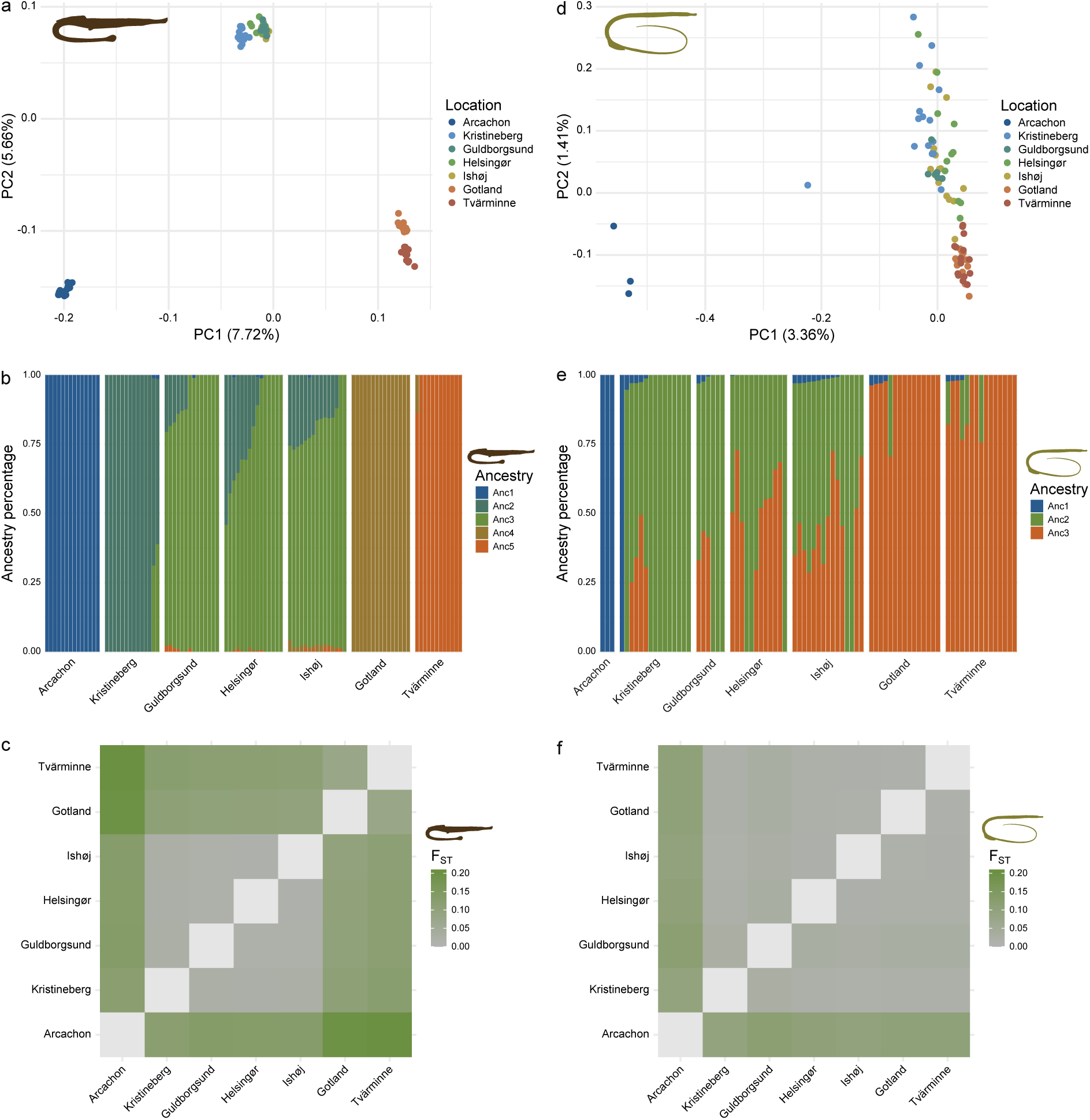
Population structure of the broadnosed pipefish (*Syngnathus typhle*) (a, b, c) based on 154,292 LD-pruned SNPs, and the straightnose pipefish (*Nerophis ophidion*) (d, e, f) based on 1,504,488 LD-pruned SNPs. (a) PCA plot for individuals of *S. typhle*. Each dot represents an individual, which is colored by sampling location. (b) Admixture plot for individuals of *S. typhle* based on genotype likelihoods. Each bar represents an individual, which are grouped by sampling site and colors correspond to the k=5 ancestral clusters. Plots for other k values can be found in Figure S4. (c) Pairwise F_ST_ for *S. typhle* based on allele frequencies per population. (d) Same plot as (a) for individuals of *N. ophidion*. (e) Same plot as (b) but for individuals of *N. ophidion* for k=3 ancestral clusters. Results for different k values are shown in Figure S4. (f) Same plot as (c) but for *N. ophidion*.

In comparison, *N. ophidion* showed weaker population structure. The Atlantic population (Arcachon) formed a distinct cluster in the PCA, while all other populations formed a gradient from west to east (Figure 2d). The PCs also explained less variance than in *S. typhle* with 3.36% for PC1 and 1.41% for PC2, and PC3 did not resolve any obvious clustering based on locations (Figure S3b). Admixture analysis on the other hand favored three ancestry proportions, one primarily in the Atlantic (Arcachon), one in the Danish Straits (Kristineberg, Guldborgsund, Helsingør, Ishøj), and a third cluster in the Baltic Sea (Gotland, Tvärminne) (Figure 2e). There was evidence of multiple individuals of mixed ancestry, especially between the clusters of the Danish Straits and the Baltic Sea, which matches the pattern seen in the PCA of less clear separation between the clusters. Interestingly, one individual sampled in Kristineberg (Danish Straits) showed primarily Atlantic ancestry. In the PCA plot, this individual also appeared in between the other individuals from Kristineberg and those from Arcachon (Figure 2d). Weaker population structure compared to *S. typhle* was also evident in F_ST_, comparisons (Figure 2f), where a separation was only clearly visible, but lower, between the Atlantic and all other populations (F_ST_=0.09-0.10), while differentiation was very low between the Baltic Sea populations and the Danish Straits (F_ST_=0.01-0.03).

### Standing genetic diversity is lower in the eastern Baltic Sea

We estimated levels of genetic diversity in each population using two metrics (Figure 3). Observed heterozygosity for each individual were similar on average between the species, with higher variability in *S. typhle* (mean *H*_O_=0.27, SD=0.04) than in *N. ophidion* (mean *H*_O_=0.28, SD=0.01, Figure 3a). At the population level, observed heterozygosity was overall relatively similar in the Atlantic and Danish Strait populations (range of means *S. typhle*: *H*_O_=0.28-0.30, range of means *N. ophidion*: 0.28-0.29), but notably lower in the Baltic populations in *S. typhle* (range of means: *H*_O_=0.21-0.22) and slightly lower in *N. ophidion* (both populations with the same mean: *H*_O_=0.27, Figure 3a). Nucleotide diversity estimated over windows of the genomes of both species revealed lower average and less variable values in *S. typhle* (mean π=0.001, SD=0.001) than in *N. ophidion* (mean π=0.003, SD=0.002, Figure 3b). At the population level, the Baltic populations of *S. typhle* had lower average nucleotide diversity (both populations with same mean: π=0.0009) than the remaining populations (range of means: π=0.0011-0.0012), while *N. ophidion* showed little difference between populations (Figure 3b).

**Figure 3:**
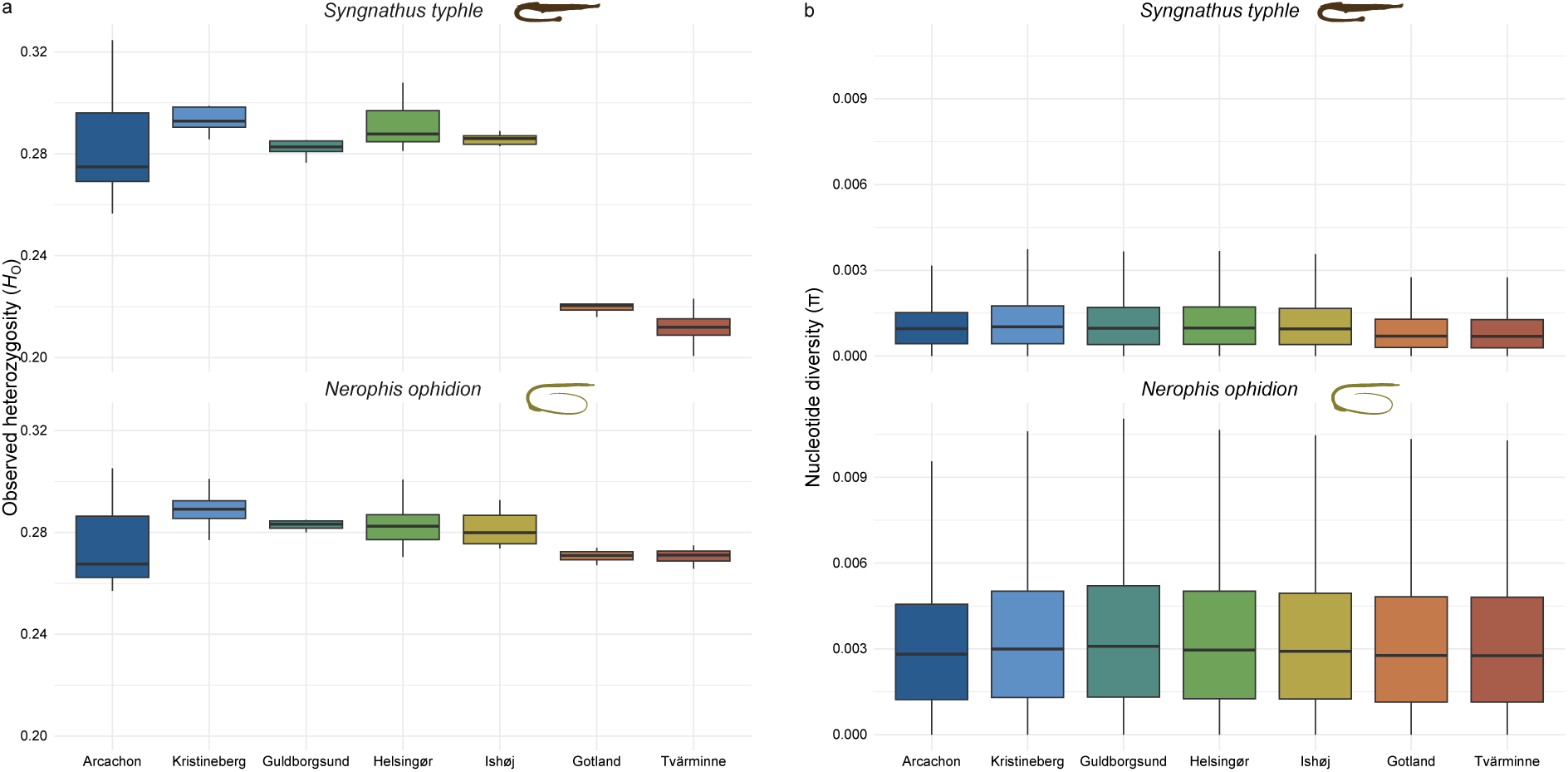
Genetic diversity of the broadnosed pipefish (*Syngnathus typhle*) based on 1,488,252 SNPs, and the straightnose pipefish (*Nerophis ophidion*) based on 19,565,453 SNPs. (a) Individual-level observed heterozygosity (*H*_O_) per population for the two species. (b) Nucleotide diversity (π) across 50 kb sliding windows with a 10 kb step size of the genome per population for the two species.

### Identifying molecular convergence in the eastern Baltic Sea populations

To investigate whether populations in the Baltic Sea experienced similar molecular changes in the two species, we evaluated convergence at four levels of genomic organization: chromosomes, pathways, genes, and individual nucleotides (Figure 4). The seven locations were grouped according to the population structure identified above into Atlantic (Arcachon), Danish Straits (Kristineberg, Guldborgsund, Helsingør, Ishøj) and Baltic Sea (Gotland, Tvärminne) (Figure 2). Because the Baltic Sea represents the focal environmental transition, all analyses compared the Baltic Sea populations with Atlantic and Danish Straits populations. With this conservative approach of requiring convergence candidates to be present not only in the Atlantic-Baltic Sea comparisons but also in Danish Straits-Baltic Sea comparisons we simultaneously address the limited sample size in the Atlantic consisting of a single location and only three samples for *N. ophidion*. Therefore, we first identified genomic regions showing signatures of divergence in the Baltic Sea populations within each species and then assessed whether these patterns converged across species at the chromosome, pathway, gene, and nucleotide level (Figure 4).

**Figure 4:**
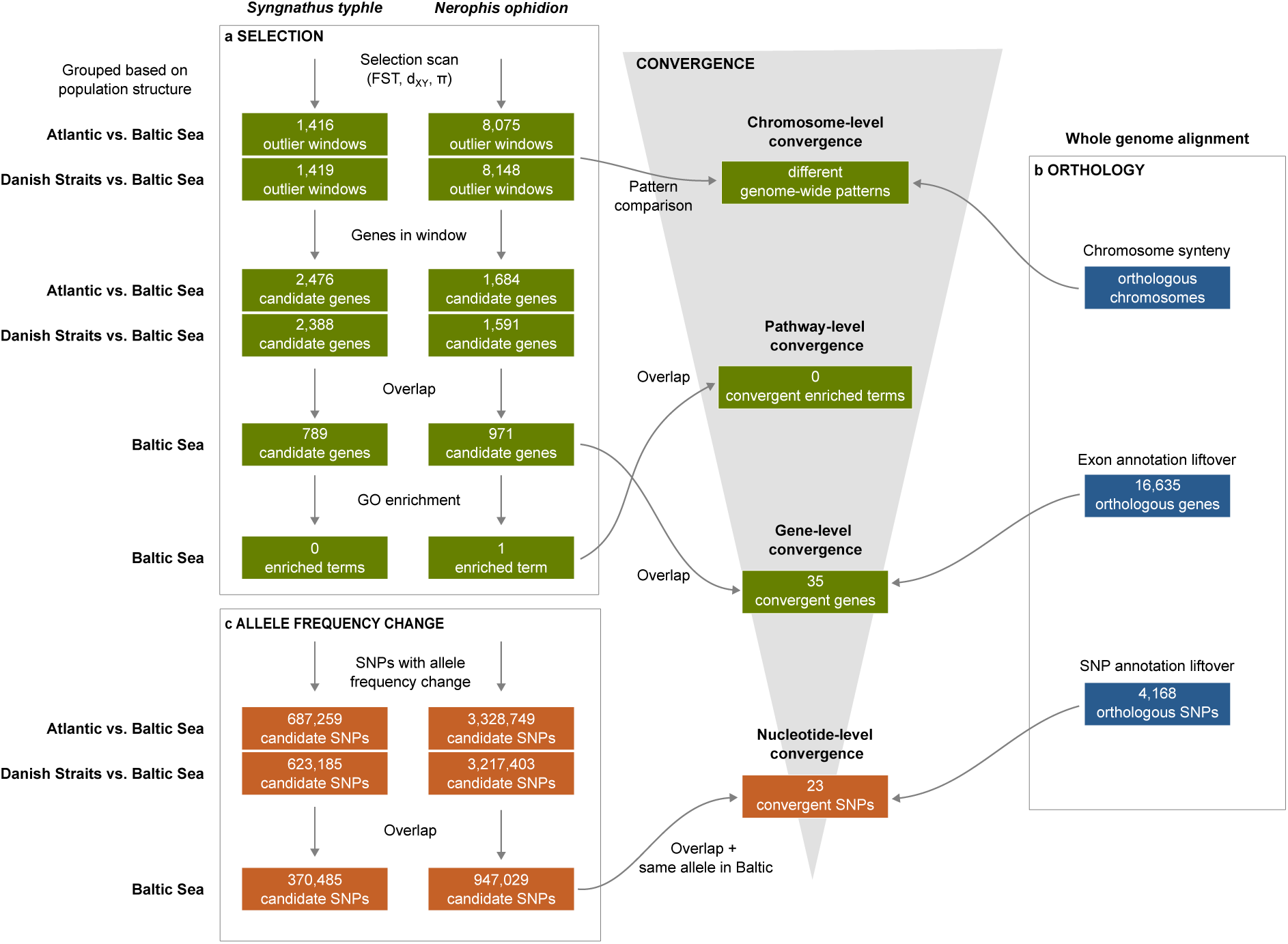
Workflow on how convergence was identified at the different molecular levels (grey arrow) by including information based on (a) selection scans, (b) orthology, and (c) allele frequency changes.

### Species-specific genomic architectures in the Baltic Sea

To assess potential convergence on the chromosome level, we compared the distribution of population differentiation (F_ST_), absolute population divergence (d_XY_) and nucleotide diversity (π) across chromosomes in the two species (Figure 4a). Genome-wide patterns of differentiation were congruent between the Atlantic-Baltic Sea and Danish Straits-Baltic Sea comparisons (Figure S5), and we therefore focus on the latter. Consistent with the findings from population structure analysis (Figure 2c, Figure 2f), genome-wide differentiation was higher in *S. typhle* (mean F_ST_=0.15) than in *N. ophidion* (mean F_ST_=0.03, Figure 5). However, the chromosomal distribution of differentiation differed markedly between species. Whereas F_ST_ outlier windows were distributed evenly across chromosomes of *S. typhle* (Figure 5a), differentiation in *N. ophidion* was concentrated on a few chromosomes (Figure 5b).

**Figure 5:**
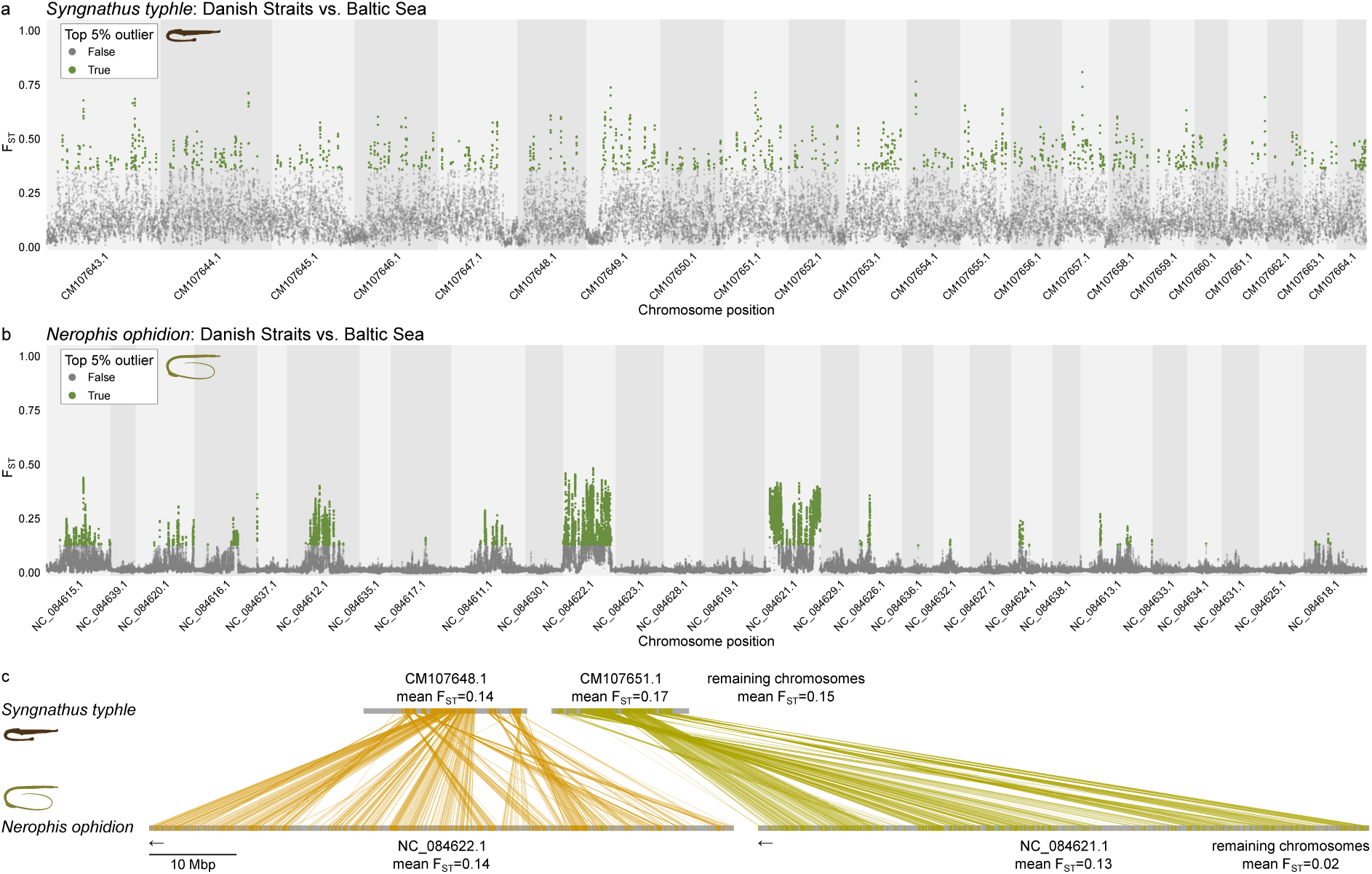
Chromosome-level differentiation of the broadnosed pipefish (*Syngnathus typhle*) and the straightnose pipefish (*Nerophis ophidion*) using F_ST_ windows across the whole genome. (a) Manhattan plot showing F_ST_ values across the whole genome between Danish Straits and Baltic Sea populations of *S. typhle*. The top 5% outliers are colored in green. (b) Same plot as (a) for *N. ophidion*. Chromosomes are ordered according to their syntenic relationships with *S. typhle* (synteny of all chromosomes shown in Figure S7). (c) Synteny of chromosomes NC_084621.1 and NC_084622.1 of *N. ophidion* with elevated overall F_ST_ values and the corresponding chromosomes CM107648.1 and CM107651.1 of *S. typhle*. Average F_ST_ values are given per visualized chromosome and for all remaining chromosomes. Reverse complemented sequences in *N. ophidion* are indicated with arrows.

In *N. ophidion*, two chromosomes (NC_084621.1 and NC_084622.1), contained extensive clusters of F_ST_ outlier windows throughout (Figure 5b, Figure S5) and had much higher differentiation (NC_084621.1 mean F_ST_=0.13 and NC_084622.1 mean F_ST_=0.14) than the remaining chromosomes (mean F_ST_=0.02) (Figure 5c, Figure S6). Elevated differentiation in these chromosomes was not associated with unusual sequencing depth or mapping rates relative to the other chromosomes. This pattern was present in both Baltic Sea populations (Gotland and Tvärminne) of *N. ophidion*, while comparisons between these two populations showed no elevated differentiation from other chromosomes (Figure S8). In addition, alleles present in the Danish Straits reference genome occurred at consistently lower frequency on both chromosomes in Baltic Sea populations (Figure S9) indicating a chromosome-wide shift in allele frequencies. Chromosome NC_084621.1 showed d_XY_ values towards the upper end of the genomic distribution together with slightly lower π in Baltic Sea populations, while NC_084622.1 had low d_XY_ compared to other chromosomes and notably reduced π in Baltic Sea populations (Figure S10, Figure S11, Figure S12). Together, these results indicate that the chromosome-scale differentiation in *N. ophidion* involves broad allele frequency shifts in both chromosomes despite differences in absolute divergence and diversity.

Synteny analysis based on the whole genome alignment (Figure 4b) showed that these chromosomes are orthologous to chromosomes CM107648.1 and CM107651.1 of *S. typhle* (Figure 5c, Figure S7). In *S. typhle*, CM107651.1 has slightly elevated chromosome-wide differentiation (mean F_ST_=0.17) compared to the remaining chromosomes (mean F_ST_=0.15), indicating that this chromosome contains windows of greater than average differentiation, although nowhere as pronounced as in the orthologous chromosome of *N. ophidion*. Overall, the contrasting patterns indicate distinct genomic architectures of adaptation in the two species with only limited evidence for chromosome-level convergence on one of the chromosomes.

### No pathway-level convergence but shared candidate genes associated with Baltic Sea divergence

For putative pathway-level convergence, genes located within windows of high F_ST_ were tested for gene ontology (GO) term enrichment within each species and enriched categories were compared across species (Figure 4a). The 789 outlier genes in *S. typhle* were not enriched in any GO terms, and the 971 outlier genes in *N. ophidion* were enriched in one GO term (ubiquitin-like protein conjugating enzyme binding). Hence, no GO term enrichment was shared between the two species. We also found no enrichment of GO terms on the genes located on the outlier chromosomes.

Despite the absence of shared enriched pathways, we identified evidence for convergence at the gene level. We focused on orthologous genes identified from a whole genome alignment between the two species (Figure 4b), which were located within outlier F_ST_ windows in both comparisons with the Baltic Sea and both species (overlap highlighted in Figure S13). This approach identified a set of 35 orthologous genes repeatedly associated with Baltic Sea divergence (Table 1, Table S4), of which 32 were located on the two highly differentiated chromosomes of *N. ophidion* (24 genes on NC_084621.1 and 8 genes on NC_084622.1). Based on available genome annotations and functional annotation with eggNOG-mapper, gene names could be assigned to 24 of these genes, while the remaining genes lacked informative annotations, including one gene without description. Although no GO term enrichment was detected, the candidate genes were associated with functions including lipid and energy metabolism (e.g. the apolipoprotein cluster, *bckdhb*, *ldha*, *slc2a4rg*), signal transduction (*cblc*, *hrasb*, G-protein coupled receptors), transcriptional regulation (zinc finger proteins), immune function (*bcl3, ifnlr1, tsg101*), protein and RNA processing (*cgh-1, man1c1*), and development (*enc, iglon5, lrrc56, myom3, rassf7a*) (Table 1, Table S4).

**Table 1:**
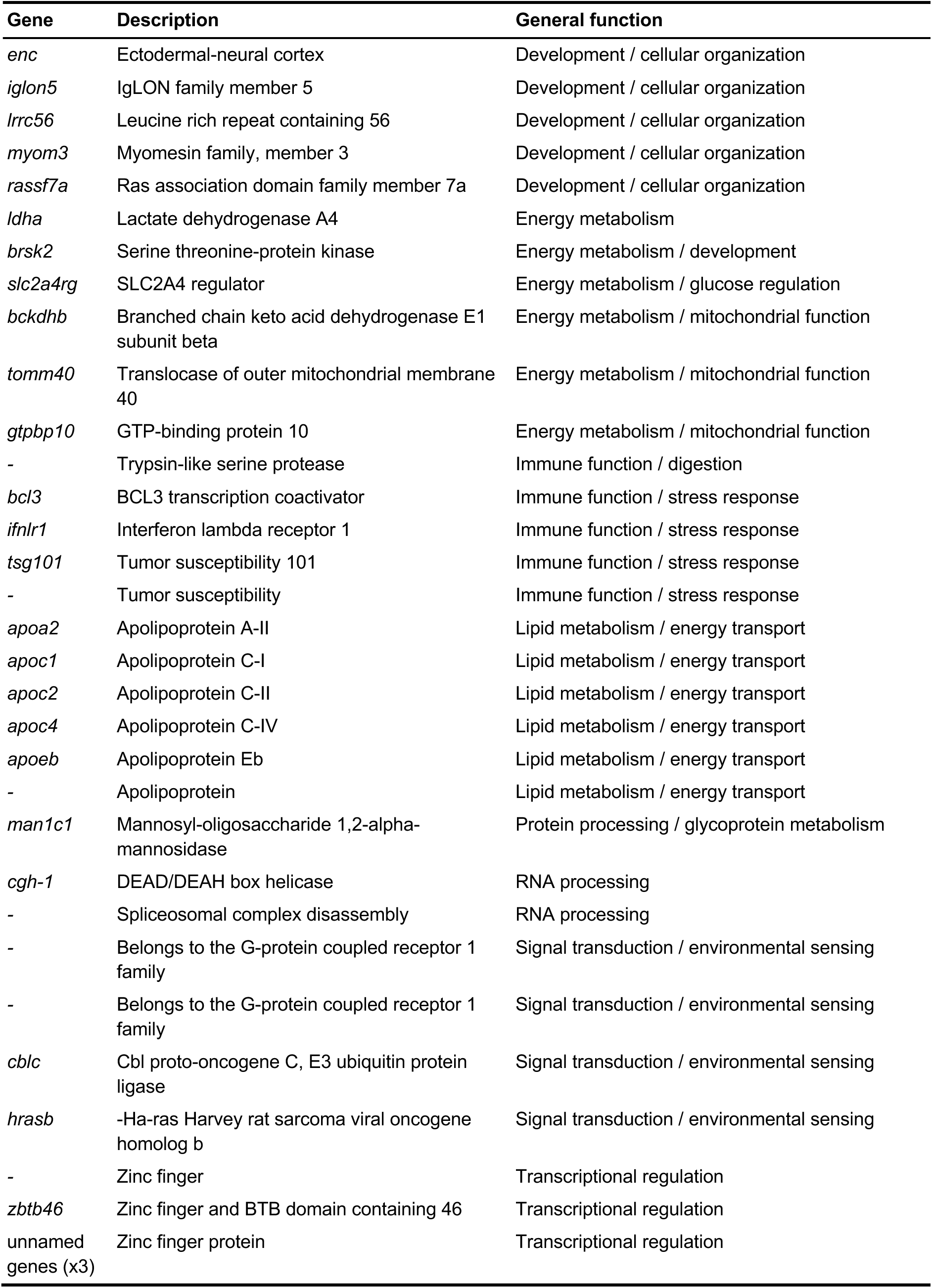
List of putatively convergent genes. Where possible, gene names and descriptions were summarized for both species. Functions were grouped using high level GO-terms. A dash (−) in the gene name indicates that the gene has no name assigned by eggNOG-mapper. Detailed information on the gene annotations can be found in Table S4.

Within the 35 convergent candidate genes, five SNPs occurred at orthologous positions in both species (Table S5). These variants were located in *slc2a4rg* (N=2), *bcl3*, *lrrc56* and a gene related to spliceosomal complex disassembly. However, all five SNPs were located outside coding regions and did not necessarily show changes in allele frequency that are specific to the Baltic. We therefore performed a dedicated nucleotide-level analysis to identify convergent SNPs irrespective of their occurrence within large differentiated genomic windows.

### Convergent allele-frequency shifts at orthologous SNPs indicate nucleotide-level convergence

To test whether convergence extended below the level of genes to the nucleotide level, we searched for orthologous SNPs from the whole genome alignment (Figure 4b) that showed parallel allele frequency shifts (>10%) toward the Baltic in both species (Figure 4c). This means that what is considered the major allele in the Baltic occurred at lower frequency in Atlantic and Danish Strait populations or that there is a different major allele in the Baltic than in the Atlantic and Danish Strait populations. This resulted in a set of 86 candidate SNPs. To identify a more stringent set of candidates of convergence, we additionally required both species to share the same major allele in the Baltic populations. This analysis resulted in a total of 23 putatively convergent SNPs (Figure 4, Figure 6).

**Figure 6:**
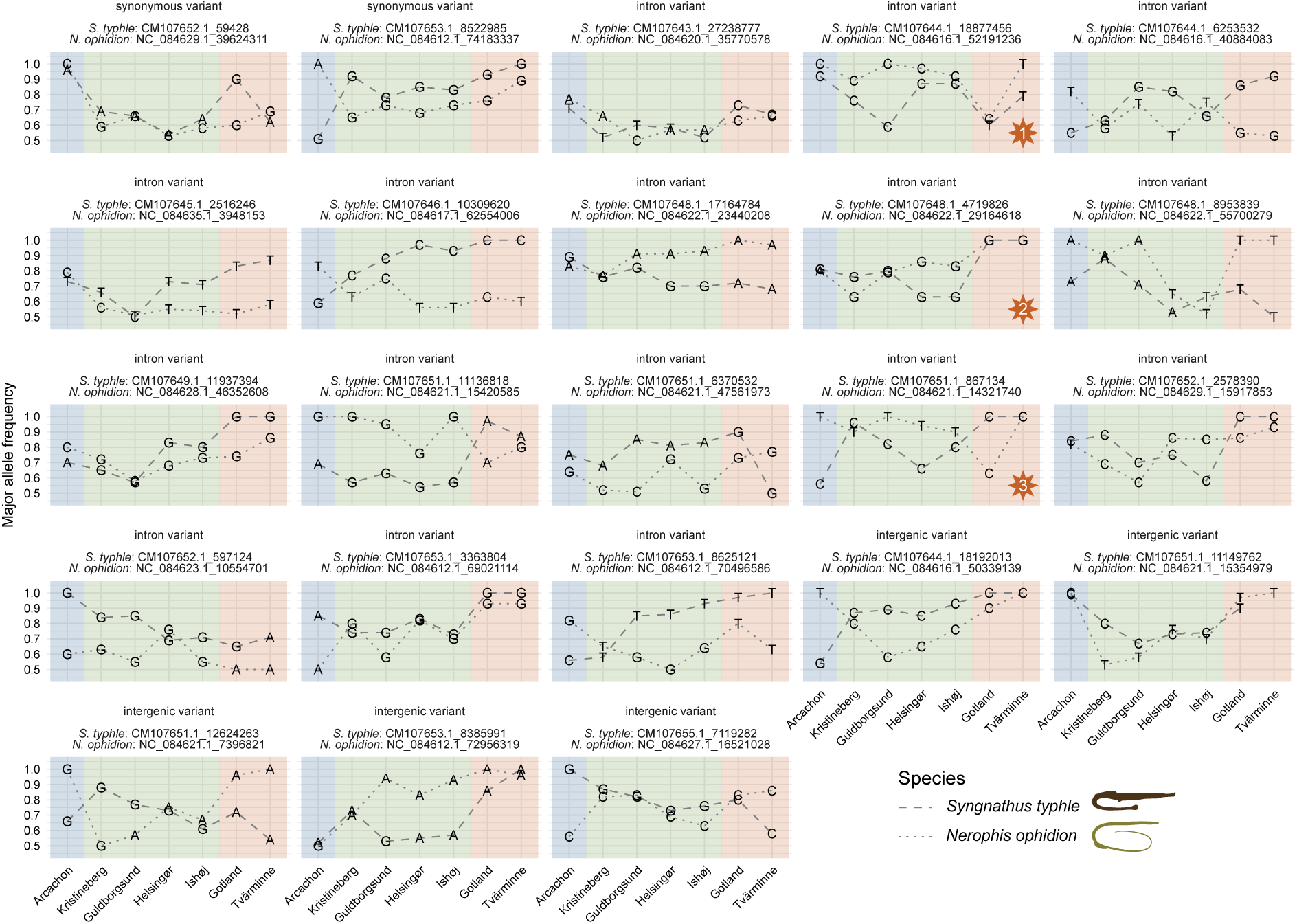
Putative convergent SNPs of the broadnosed pipefish (*Syngnathus typhle*) and the straightnose pipefish (*Nerophis ophidion*). The panels show allele frequencies of the major allele for each population for each of the 23 convergent variants for which (i) both species show the same major allele in the Baltic Sea populations (highlighted in red), and (ii) the major allele of the Baltic Sea populations is either different to the major allele in the Atlantic (blue) and to the major allele in the Danish Straits (green), or (iii) there was an allele frequency change of at least 0.1 in the Baltic Sea compared to the other two groups. Genomic coordinates and genomic location are given on top of each panel. Three SNPs representing distinct patterns used as examples in the text are highlighted with an orange star.

Of these SNPs, two were synonymous variants located within coding regions, 16 were located within introns, and the remaining 5 were outside genes (Figure 6). Despite meeting the same convergence criteria, these SNPs exhibited diverse patterns of allele frequency change across populations. At the position of some SNPs, both species showed a shift from one major allele in Atlantic and Danish Straits populations to a shared major allele in the Baltic Sea (Figure 6, example 1). At other loci, the frequency of the Baltic-associated allele increased progressively towards the Baltic, reaching full fixation of the same allele in both species (Figure 6, example 2). Other SNPs showed different patterns in the two species, with fixation of the Baltic-associated allele in one species but not the other, while still resulting in the same major allele being favored in Baltic populations of both species (Figure 6, example 3).

## Discussion

Here, we generated a new reference genome for *S. typhle* and analyzed whole-genome resequencing data for the pipefish species *S. typhle* and *N. ophidion*, sampled along the Atlantic-Baltic environmental gradient to investigate the genomic consequences of their independent colonization of the Baltic Sea. Although population structure in *S. typhle* has previously been investigated using few genetic markers and reduced representation sequencing (Wilson and Eigenmann Veraguth 2010; Goehlich et al. 2021; Knutsen et al. 2022), this study provides the first whole-genome assessment in this species. For *N. ophidion*, this represents the first population genetic analysis and therefore the first insight into the evolutionary consequences of the Baltic Sea colonization in this species. Both species exhibited elevated population differentiation and reduced genetic diversity in Baltic Sea populations, consistent with shared demographic effects associated with the Baltic Sea colonization. Colonization of the Baltic Sea did not result in the same patterns at the level of genomic architecture, because the two species used markedly different chromosomal configurations of divergence. In *S. typhle*, signals of selection were distributed across much of their compact genome, whereas in *N. ophidion* they were concentrated on a small number of chromosomes, suggesting that adaptation involved different genomic regions in the two species. Nevertheless, we identified several orthologous genes and SNPs showing parallel signals of differentiation between Baltic and other populations. Together, these findings reveal a scale-dependent convergence with genomic architectures of adaptation differing between species, whereas convergence emerged at the level of genes and nucleotide variants.

### Population structure and diversity are congruent following Baltic Sea colonization

Despite similar ecologies and expected dispersal limitations, *S. typhle* and *N. ophidion* differed markedly in their population differentiation, while the geographic patterns were broadly similar. Genetic differentiation among populations of *S. typhle* was strong, whereas *N. ophidion* exhibited very weak differentiation from the Danish Straits to Finland, as evidenced by PCA and F_ST_ analyses. Nevertheless, both species exhibited distinct ancestry components associated with the Baltic Sea. This places the two pipefish species among the many marine taxa that form genetically distinct populations in the Baltic (Wennerström et al. 2017; Leder et al. 2021; Geburzi et al. 2022; Andersson et al. 2023). At the same time, the contrasting levels of differentiation between *S. typhle* and *N. ophidion* add evidence that the strength of population structure across the Danish Strait-Baltic gradient is highly species-specific (Wennerström et al. 2013). Together, these patterns suggest that colonization of the Baltic Sea repeatedly promoted population differentiation, while species-specific characteristics such as dispersal potential likely shape its extent.

Strong structuring in *S. typhle* is consistent with previous studies (Wilson and Eigenmann Veraguth 2010; Knutsen et al. 2022) and expected given the generally low dispersal potential of pipefishes. In contrast, the weak structure and extensive admixture observed in *N. ophidion* suggest greater connectivity among populations. It is possible that there are differences in larval dispersal abilities, with the small juveniles of *N. ophidion* possibly dispersing more effectively than the larger *S. typhle* juveniles (Braga Goncalves et al. 2016). It is also possible that *N. ophidion* juveniles and adults raft more than *S. typhle* on detached seagrass, a behavior that has been described for other syngnathids with grasping abilities (Bertola et al. 2020; Li et al. 2021). Additional support for broad connectivity comes from a Kristineberg individual that occupied an intermediate position between the Danish Strait and the Atlantic clusters and showed predominantly Atlantic ancestry. Given the large sampling gap between Kristineberg and Arcachon, this individual may stem from an unsampled intermediate population along the Atlantic coast from which it dispersed to Kristineberg.

Genetic diversity was lower in Baltic Sea populations than in Atlantic and Danish Straits populations of both species, with the reduction being particularly pronounced in *S. typhle*. Reduced genetic diversity is a common feature of Baltic Sea populations of different species (Johannesson and André 2006; Wennerström et al. 2017). The lower diversity observed here may reflect founder effects associated with the initial colonization of the Baltic Sea. Contemporary demographic processes may also contribute as effective population sizes of Baltic populations are estimated to often be very small (*N*_e_ < 50, (Wennerström et al. 2017)), which can accelerate loss of genetic diversity through genetic drift. Even though adaptation is likely to proceed faster from standing genetic variation than adaptation relying on new mutations (Barrett and Schluter 2008), reduced diversity does not necessarily imply reduced adaptive potential (Liu et al. 2019; Abson et al. 2026). The reduced diversity observed in Baltic populations of pipefishes did not seem to hinder adaptive potential, as both species showed signals of selection and evidence of convergence.

### Molecular convergence is weak at broader genomic levels but evident at more fine-scale molecular scales

Convergence between Baltic Sea populations of the two species was investigated at chromosome, pathway, gene, and nucleotide level. By performing comparisons of different populations, we are able to identify convergent signals that would be hidden in a comparison at the species level. At the chromosome level, the genomes of the species showed different patterns of differentiation when comparing the Baltic Sea to the Atlantic and the Danish Straits. For *N. ophidion* differentiation was concentrated on two chromosomes that were highly differentiated across most of their length, accompanied by chromosome-wide shifts in allele frequencies in Baltic Sea populations. In contrast, signals of differentiation in *S. typhle* were distributed across chromosomes, although we find that the orthologous chromosome to one of the *N. ophidion* outlier chromosomes (NC_084621.1) had higher mean chromosome-wide differentiation than the other chromosomes. This indicates that in both species, genes on these chromosomes are under stronger selection than the rest of the genome.

The contrasting genomic architectures observed in the two species may reflect differences in population connectivity. Theory predicts that when gene flow is limited, adaptive divergence can accumulate across loci distributed throughout the genome (as in *S. typhle*), whereas higher gene flow can produce fewer, larger-effect, and more tightly linked adaptive loci (as in *N. ophidion*) because linkage helps maintain locally adapted allele combinations despite gene flow (Yeaman and Whitlock 2011). It is possible that large structural rearrangements are involved in the differentiation of the two *N. ophidion* chromosomes in the Baltic Sea populations, similar to large chromosomal inversions identified in different populations of the long-snouted seahorse (Meyer et al. 2024). Such inversions or other structural rearrangements suppress recombination and maintain adaptive haplotypes in the face of gene flow that can facilitate local adaptation (Kirkpatrick and Barton 2006; Yeaman 2013; Wellenreuther and Bernatchez 2018; Wonneberger et al. 2023). For example, in the Atlantic silverside, such regions of reduced recombination are suggested to facilitate adaptation under divergent selection despite ongoing gene flow between populations (Akopyan et al. 2025).

The two differentiated chromosomes in *N. ophidion* showed different patterns of diversity and divergence, suggesting that different evolutionary processes may underlie their differentiation. Chromosome NC_084621.1 had somewhat elevated nucleotide divergence relative to the other chromosomes, consistent with a scenario of ancient divergence with ancestral polymorphism of the differentiated chromosome (Ma et al. 2018; Akopyan et al. 2025; Luzuriaga-Aveiga et al. 2025), while chromosome NC_084622.1 showed reduced nucleotide diversity in the Baltic Sea populations, a pattern more consistent with a selective sweep (Moinet et al. 2022; Akopyan et al. 2025; Luzuriaga-Aveiga et al. 2025). These patterns may reflect a combination of selection, reduced recombination, and genetic diversity changes within populations (Charlesworth 1998; Cruickshank and Hahn 2014; Burri 2017), although analyses of structural variation and recombination landscapes will be required to distinguish among these possibilities.

We found no evidence for convergence at the pathway level. Pathway-level convergence has for example been shown in bats (Morales et al. 2024), and it has been suggested that it may be more common than gene-level convergence when divergence time is large (Hoitinga and Birkeland 2025). While limited statistical power and incomplete gene annotations may contribute to the absence of pathway-level convergence in this study, it is also possible that selection repeatedly targeted a small number of genes rather than entire biological pathways, which would not be detected with our enrichment approach.

We identified 35 orthologous candidate genes associated with Baltic Sea divergence in both species. Many were linked to lipid and energy metabolism, immune defense and gene regulation, processes that are reasonably expected to be impacted by differences in salinity, temperature and resource availability between marine and brackish environments. In the Baltic, *S. typhle* is known to grow to smaller sizes (Wilson et al. 2020; Goehlich et al. 2021), possibly reflecting the increased metabolic demands of osmoregulation and other environmental stressors. Similarly, variation in prey resources, temperature, and salinity can alter energy demands in species inhabiting the Baltic Sea (Morgan and Iwama 1991; Volkoff and Rønnestad 2020; Gallagher et al. 2022), potentially explaining the repeated involvement of metabolic genes. Several shared candidate genes were also related to immune function, consistent with different pathogen communities between marine and Baltic habitats (Fleischmann et al. 2022; Mazur-Marzec et al. 2024) as well as the previously described effect of salinity stress on the immune defense in pipefish (Birrer et al. 2012; Goehlich et al. 2021). This suggests that adaptation to different immune challenges may represent another convergent response following Baltic Sea colonization.

Surprisingly, we did not identify the classical osmoregulatory genes often implicated in salinity adaptation (e.g. *slc12a*, *atp6* and *cftr*), including in freshwater pipefishes (Flanagan et al. 2021) and in other fishes crossing the Baltic Sea salinity gradient (Leder et al. 2021). This may indicate that Baltic Sea adaptation in these pipefishes is mediated through other physiological mechanisms or through changes in the regulation of genes, rather than repeated evolution of the same ion transporters. Consistent with this interpretation, several candidate genes are involved in regulatory functions, suggesting that changes in gene expression may contribute to adaptation.

For convergence at the nucleotide level, we also found potential relevance of regulatory elements based on the 23 positions where both species showed the same nucleotide at elevated frequencies in the Baltic Sea populations relative to non-Baltic populations. Although such parallel allele frequency shifts are consistent with the action of natural selection, the functional significance of these variants remains unclear. None of these convergent SNPs cause amino acid substitutions, frameshifts, or chances to start or stop codons, suggesting that selection is unlikely to be driven by repeated changes to protein-coding sequences in this system. Instead, the predominance of SNPs in introns and intergenic regions points towards a potential role in gene regulation (e.g. splicing or gene expression). Given the ca. 58 million years of divergence between the two species and their substantial differences in genome size and chromosomal organization, it is possible that orthologous SNPs could occur in divergent genomic contexts in the two species and therefore may not retain the same functional roles in both species. Future work could focus on identifying whether these putatively convergent SNPs lie in enhancer regions or at transcription factor binding sites in order to clarify whether they contributed to adaptation to the Baltic Sea environment.

## Future directions

Overall, our findings support the view that convergence strongly depends on the biological level examined (Greenway et al. 2024). Future work should focus on identifying the causal mechanisms underlying the patterns observed here, particularly through analyses of structural variation, recombination landscapes, and pangenomes instead of relying on a single reference genome (Secomandi et al. 2025) in order to assess the extent to which structural variations contribute to population divergence across the Baltic Sea gradient. Functional validation of candidate genes and SNPs using transcriptomic (e.g. RNA-seq, qPCR) and experimental approaches, together with improved annotation of regulatory regions (e.g. via ATAC-seq, ChIP-seq or bioinformatic approaches) will help determine whether convergent adaptation is mediated through changes in gene regulation. Finally, integration of genomic, environmental, and phenotypic information as well as expanded sampling will improve our understanding of adaptation along environmental gradients and the relative roles of genetic adaptation and phenotypic plasticity (Goehlich et al. 2021).

## Conclusion

Taken together, our results reveal a scale-dependent pattern of convergence. Despite experiencing the same environmental transition, *S. typhle* and *N. ophidion* evolved markedly different genomic architectures in the Baltic, although one orthologous chromosome appeared to be under stronger selection in both species, and showed no evidence for convergence at the pathway level. Nevertheless, the detection of convergent genes and nucleotides suggests that common selective pressures can still produce shared molecular responses despite considerable evolutionary divergence. These findings suggest that connectivity, genetic diversity, and genome organization can shape how adaptation is distributed across the genome, although some targets may ultimately be favored by selection. More broadly, this study demonstrates how whole genome alignments and comparative analysis across populations can bridge population-level processes and macroevolutionary time scales, allowing consequences of a recent environmental transition to be compared across lineages that diverged approximately 58 million years ago.

## Methods

### Diploid genome assembly for *Syngnathus typhle*

Two individuals of broadnosed pipefish (*S. typhle*) were collected snorkeling over eelgrass with a net in Køge Bay, Denmark on 14 September 2022 (individual codes Syntyp1 and Syntyp11). A physical voucher of individual Syntyp11 was deposited at the Natural History Museum of Denmark’s (catalog number NHMD, ZMUC P2398900). The specimens were collected under a permit (21-450C) to the Natural History Museum Denmark. The animals were killed using MS-222 after which tissue samples were flash frozen in liquid nitrogen.

The reference genome was built from tissues collected from individual Syntyp11. A liver sample was used for PacBio HiFi sequencing at the Yggdrasil Eukaryotic Reference Genome Facility at the University of Copenhagen as described in (Kliver et al. 2025). High molecular weight (HMW) DNA was extracted from 21 mg liver using MagAttract HMW DNA Kit (Qiagen) and following the manufacturer’s protocol. One PacBio 8M wells SMRT cell was loaded and sequenced on a Pacific Biosciences SEQUEL IIe platform.

A chromatin-capture library was built using the Arima High Coverage HiC kit (Arima Genomics, San Diego, CA, USA) from a 20mg sample of brain and eyes. Sequencing of HiC libraries was sent to Novogene and done on an Illumina NovaSeq 6000 platform using PE150 bp chemistry aiming for 100 Gb of data.

To assist genome annotation, RNA library preparation was performed for two tissues (4mg of eyes, 6mg of gills) collected from individual Syntyp1. RNA was extracted using the Quick-RNA Miniprep Plus Kit (Zymo Research), following the manufacturer’s instructions. RNA integrity and concentration were assessed using RNA 6000 Nano kit with a BioAnalyzer (Agilent Technologies) and a Qubit Fluorometer (Thermo Fisher Scientific, Waltham, MA, USA), using the Invitrogen Qubit RNA Broad Range assay. RNA samples had RIN values of 7.7 (eyes) and 6.2 (gills). RNA libraries were constructed using Revelo mRNA-Seq for MagicPrep NGS (Tecan) with 500 ng of RNA as input on a MagPrep machine (Tecan). It involved polyA selection and both strands synthesis. Library fragment size distribution and concentration were assessed using High Sensitivity DNA kit with a BioAnalyzer (Agilent Technologies) and a Qubit Fluorometer (Thermo Fisher Scientific, Waltham, MA, USA) using Qubit DNA High Sensitivity assay, respectively. RNA-seq libraries were sent to Novogene for sequencing on an Illumina NovaSeq 6000 using PE150 bp chemistry aiming for 30Gb of data per library.

The diploid genome was assembled using the AssemblyBrute v0.1 pipeline (Kliver 2023) following the VGP assembly approach version 2 (Larivière et al. 2024) with additional quality control steps, as described in (Kliver et al. 2025). Genome assembly statistics were obtained with ragtag v2.0.1 (Alonge et al. 2022). Genome completeness was assessed with BUSCO v5.8.0 (Manni et al. 2021) using the actinopterygii_odb10 database and miniprot (Li 2023) as the gene predictor.

Gene annotation for haplotype 1 was generated as part of the VGP phase 1 using EGAPx (NCBI 2024), which employed an external version of the NCBI RefSeq annotation pipeline (Formenti et al. 2026). This annotation was transferred to haplotype 2 using liftoff version 1.6.3 (Shumate and Salzberg 2021). Missing phase information was added with agat_sp_fix_cds_phases.pl from agat v1.4.0 (Dainat et al. 2026).

### Population-genetic sampling

A total of 180 individuals from the two pipefish species *N. ophidion* and *S. typhle* were collected from seven different locations across Europe (France, Sweden, Denmark to Finland, Figure 1a). One additional individual of *S. typhle* was not included after initial analysis because it showed unusually high relatedness with all individuals, likely stemming from contamination (Figure S1).

Permits for sampling and handling the animals were in place where necessary according to national regulation. Collection permits were as follows: Arcachon (using permits from the Station Marine d’Arcachon); Denmark (21-450C); Gotland and Kristineberg (Idnr 004797, Dnr 5.8.18-04405/2023); Tvärminne (using permits from the Tvärminne Zoological Station). Samples were imported to Denmark using an import permit (DK-3-oth-710475, J.nr. 2025-61261, Ref. LHEN 2024-12-7186-03365, 2024-12-711-07957). Individuals from Guldborgsund, Denmark were sampled in 2022, while the remaining locations were sampled in summer 2025, either by snorkeling with a hand net, by scuba diving or with a beam trawl pulled behind a boat (Table S1, Table S2). The individuals were euthanized with MS-222 (FELASA certificate number ABD-F032/10/25-211 and additionally for Swedish sampling locations: completed course on Swedish legislation & Ethics, animal welfare, and 3R). Muscle tissue was taken for DNA extraction. Samples were stored in the freezer at −20°C or −70°C until processing.

### DNA extraction, library preparation, and sequencing

For each individual, DNA was extracted from muscle tissue using Qiagen DNeasy Blood & Tissue Kit under standard protocol. DNA concentration was quantified using a Qubit 1X dsDNA High Sensitivity assay on a Qubit 4 Fluorometer. Libraries for sequencing were prepared using the Illumina DNA PCR-Free Prep kit following standard protocol. Illumina DNA/RNA UD Indexes Set A and B were used. The concentration of the library was analyzed with Qubit using the ssDNA kit. Based on this concentration, the samples were pooled, adjusting the added volume to aim for equal molarity within species in order to achieve an equal sequencing coverage while adjusting for genome size differences between species. Samples were sequenced on 3 lanes of the NovaSeq X Plus (PE150) with Novogene Europe.

### Processing of sequence reads

Raw BCL data were demultiplexed with bcl2fastq v2.20.0.422 (Illumina 2017). Adapter sequence trimming and low-quality base filtering was performed using fastp v0.23.2 (Chen et al. 2018). Sliding-window trimming was applied from the 5’ front to the 3’ tail of the reads using a sliding window of 4 bases and a mean quality threshold of 15. Additional stricter filtering was applied on the 5’ end with a window size of 1 base and a mean quality threshold of 10. Sequencing quality was assessed using fastQC v0.12.1 (Andrews 2010).

Trimmed reads were mapped to the newly generated genome for *S. typhle* (haplotype 2, GCA_048301605.1) and to an available chromosome-level genome for *N. ophidion* (GCF_033978795.1, (Ramesh et al. 2024)) using bwa-mem2 mem v2.2.1 (Vasimuddin et al. 2019). The SAM output files were converted to BAM format using samtools v1.20 view (Danecek et al. 2021). After sorting the bam files with samtools sort, we used MarkDuplicates from picard-tools v3.1.0 (Broad Institute 2018) and removed overlap of paired reads with bam clipOverlap from bamutil v1.0.14 (Jun et al. 2015). Mapping quality was assessed using samtools flagstat and average coverage was calculated using picard CollectWgsMetrics (Broad Institute 2018). Finally, BAM files were filtered to only include reads from chromosomes, not unplaced scaffolds, using samtools view and indexed with samtools index.

### Variant identification, filtering, and functional annotation

Analyses were performed based on genotype likelihoods which can be advantageous over calling of genotypes when sequencing coverage varies between individuals. For each species, ANGSD v0.940 (Korneliussen et al. 2014) was run using the GATK model (McKenna et al. 2010). To ensure high read and mapping quality, flagged duplicated or overlapping reads were excluded as described above and we further only kept reads that match to exactly one position and reads where both read pairs mapped properly. Base alignment quality was calculated, and mapping quality was adjusted by 50 for excessive mismatches. Bases with a mapping quality below 20 were discarded. Output format was set to beagle, and a bcf file was created. Allele counts were enabled requiring a minimum sequencing depth of 1 and allele frequencies were calculated with major and minor alleles defined by the genotype likelihoods (Kim et al. 2011). Only sites with a minor allele frequency of at least 0.05, with non-missingness for at least 4 individuals and a p-value below 1e-6 were retained. Resulting bcf files were converted to vcf with bcftools view v1.23.1 (Danecek et al. 2021).

Evidence for linkage disequilibrium (LD) was assessed by calling genotypes with bcftools v. 1.23.1 (Danecek et al. 2021) for all SNPs for which at least 70% of individuals had at least one read and where the genotype confidence was above 90%. The output was then used to get a list of unlinked SNPs with plink v1.9.0 (Chang et al. 2015; Purcell and Chang 2015) with 200kb windows and a default minimum r2 of 0.2 (Gaunt et al. 2007). We filtered the ANGSD dataset to keep only this LD-filtered subset of SNPs for population structure analyses. To confirm the results from the LD-filtered dataset, analyses were repeated with a dataset that was filtered to keep only one random SNP per 10kb window. This is a simpler way of filtering out linked SNPs without relying on genotypes.

One chromosome in *Nerophis ophidion* (chromosome 4, NC_084614.1) showed extremely high LD and segregated with the sex of individuals (Figure S2), possibly representing a sex-linked chromosome. Hence, it was excluded from population-level analyses for *N. ophidion*.

To annotate identified variants, the position of each SNP was overlapped with the annotation of the respective reference genome (*S. typhle* haplotype 2, *N. ophidion* GCF_033978795.1-RS_2023_12). Functional annotation was added using eggNOG-mapper v2.1.12 (Huerta-Cepas et al. 2019; Buchfink et al. 2021; Cantalapiedra et al. 2021). Regions of reference and eggNOG-mapper annotations were intersected with the SNP positions. For this study, we were only interested in annotations on a gene level, hence different annotations for multiple transcripts of the same gene were merged into one by keeping one representative name and description, but all associated GO terms. We identified whether a SNP was inside an exon of the associated gene and annotated the functional consequence of all SNPs based on their position within genes using the variant effect predictor (VEP) v115.2 (McLaren et al. 2016). Because a SNP can have different functional consequences, all consequences were retained. Because genes can overlap, a SNP can have multiple associated genes, which we kept separate.

### Population structure analyses

For each species, the LD-filtered SNPs were used to avoid inflated population structure from linked variants. A PCA was performed with PCangsd v1.36.4 (Meisner and Albrechtsen 2018) providing the ANGSD beagle output file with genotype likelihoods with a minor allele frequency threshold of at least 0.05 and 500 iterations to ensure convergence of the algorithm.

Admixture analysis was performed with NGSadmix v32 (Skotte et al. 2013) using the genotype likelihoods of unlinked SNPs in beagle format as the input. For the number of ancestral clusters k, a range between 2 and 8 was tested and the minor allele frequency was set to at least 0.05. For each k, 10 replicates were performed, which were then visualized with pong v1.5 (Behr et al. 2016) with a pairwise similarity threshold of 0.85. The replicate designated as the “major mode” was chosen for each k value. Optimal k values were chosen with evalAdmix v1.0 (Garcia-Erill and Albrechtsen 2020) as the k where the major mode shows no population-level correlated residuals (Figure S14).

Fixation indices (F_ST_) were calculated for each population pair with realSFS from ANGSD (Nielsen et al. 2012). This requires allele frequency files per population, which were created with ANGSD -doSaf 1, providing the indexed reference genome and the sample size of the population. A vcf file subset to the individuals from the population was used as the genotype likelihood input. Based on these saf files per population, realSFS was run to calculate the two-dimensional site frequency spectrum (2d-SFS) for all combinations of population pairs. We ran realSFS fst index while providing the two saf files as well as the generated sfs file for population pair and realSFS fst stats in order to calculate the global weighted F_ST_ values between the population pair.

### Genetic diversity

Heterozygosity per individual was calculated with vcftools v0.1.17 (Danecek et al. 2011) based on called genotypes via bcftools v. 1.23.1 (Danecek et al. 2021) for all SNPs where the genotype confidence was above 95%. Nucleotide diversity (π) per population was calculated for 50,000 base pair sliding windows with a step size of 10,000 bases using thetaStat do_stat from ANGSD (Nielsen et al. 2012), dividing the pairwise theta (tP) by the window size. This assumes a theta of zero for all sites that were not identified as SNPs.

### Identification of convergence

In order to identify molecular convergences between the two species, we considered four levels, namely chromosome-level, pathway-level, gene-level and nucleotide-level convergences. We used evidence from orthology, selection analyses, and allele frequency change (Figure 4). Based on the population structure results, sample sites were grouped into three clusters: Atlantic (Arcachon), Danish Straits (Guldborgsund, Helsingør, Ishøj) clustering with Skagerrak (Kristineberg), and eastern Baltic Sea (Gotland, Tvärminne). With the Baltic Sea being the focal population, convergence was analyzed for Atlantic-Baltic Sea and Danish Straits-Baltic Sea comparisons. We required convergence candidates to be present not only in the Atlantic-Baltic Sea comparisons but also in Danish Straits-Baltic Sea comparisons. This deliberately conservative approach reduces the likelihood that convergence candidates are driven by the Atlantic populations being represented by a single locality (Arcachon), with small sample size for *N. ophidion* (N=3).

### Identification of orthologous chromosomes, genes and SNPs between the two species

To be able to compare the two species, a whole genome alignment was created using progressive cactus v2.8.0 (Armstrong et al. 2020). This also included other Syngnathiformes species to bridge the phylogenetic distance between the two target species. We used halSynteny from HAL tools v3.1.4 (Hickey et al. 2013) to export syntenic blocks to psl format using a maximum anchor distance and minimum block size of 5000 bp. Synteny among the chromosomes (omitting unplaced scaffolds) of the two species was visualized with ntSynt-viz (Coombe, Warren, et al. 2025; Coombe, Kazemi, et al. 2025). SNPs identified in *S. typhle* were mapped to positions in the *N. ophidion* genome with halStats from HAL tools, providing the SNP positions in BED format. SNPs of *S. typhle* that mapped to multiple positions in the genome of *N. ophidion* were filtered out. The resulting positions were then intersected with *N. ophidion* SNPs to find a set of convergences at the nucleotide level in both species. Similarly, a gene mapping was created by lifting all *S. typhle* exons to *N. ophidion* with halStats and intersecting it with the gene annotation for *N. ophidion*. Exons of *S. typhle* were grouped together by mRNA and then by gene, while summing the number of bases that overlapped with the gene of *N. ophidion*. In order to reduce false mapping of genes based on a few bases, mappings between genes had to overlap at least 50% of the gene (Figure 4b).

### Selection scans

The goal of the selection analysis was to identify genome windows that showed F_ST_ values compared to genome-wide average, while also taking nucleotide divergence (d_XY_) between populations and nucleotide diversity (π) within populations into account. The sample locations were grouped into Atlantic (Arcachon), Danish Straits (Kristineberg, Guldborgsund, Helsingør, Ishøj) and Baltic Sea (Gotland, Tvärminne) in order to perform comparisons between Atlantic-Baltic Sea and Danish Straits-Baltic Sea. Similarly to the global F_ST_, F_ST_ was first estimated per site by following the previously described F_ST_ calculation with realSFS fst print. Windowed F_ST_ could then be calculated using the command realSFS fst stats2, using a window size of 50,000 bases and a step size of 10,000. For the same windows, π values were calculated as described in the section on genetic diversity. Per site d_XY_ calculations are based on the calcDxy.R script from the ngsPopGen toolkit (Fumagalli et al. 2014), using the reference allele frequencies as the input. The results are then averaged per window, assuming a d_XY_ of zero for sites without available allele frequencies. After combining the three statistics per window, only windows are kept where F_ST_ and d_XY_ are present and which have more than 50 sites. This is aiming to prevent windows being detected as outliers based on very few SNPs. Then, windows in the top 5% quantile of F_ST_, meaning high differentiation, were chosen as outliers. Additionally, if outliers fell in the top 50% of d_XY_ they were considered as divergent. If the window was in the bottom 25% of π for the Baltic Sea population it was marked as having low diversity within the Baltic Sea. Genome-wide patterns of F_ST_, d_XY_ and π were compared between species to assess chromosome-level convergence, taking synteny based on the whole genome alignment into account. To identify which genes and SNPs lie within windows that are candidates for selection, the window ranges were overlapped with gene and SNP positions (Figure 4a).

### Convergence of pathways under selection

In order to identify potential pathway-level convergence, we tested for GO term enrichment using a list of candidate genes identified per species. All genes that were identified as outliers based on F_ST_ in the comparison of Atlantic-Baltic Sea and Danish Straits-Baltic Sea were chosen as the input set for enrichment. GO terms were associated with genes based on the eggNOG-mapper annotation. Enrichment was tested using all genes of the corresponding species as the background set. Resulting GO terms per species were intersected to identify convergently enriched pathways.

### Convergence of genes under selection

For identifying gene-level convergence between the two species, the candidate genes under selection per species were combined with the established mapping of orthologous genes between species. For each species, the intersection of candidate genes of the two comparisons (Atlantic-Baltic and Danish Straits-Baltic) was used to address the low sampling size of the Atlantic site for *N. ophidion*, and to be able to test directionality with the focus on the Baltic Sea. The candidate genes for the Baltic Sea per species were combined using the mapping between species to identify which orthologous genes are outliers in both species. This results in a list of convergent genes (Table 1, Table S4).

### Nucleotide-level convergence using allele frequency changes

To identify candidate SNPs in the Baltic Sea, instead of relying on large selection windows, we focused on changes in allele frequency of individual SNPs when comparing Atlantic-Baltic Sea or Danish Straits-Baltic Sea populations. SNPs were considered as candidates for a specific species and one of the population comparisons if the major allele differed in the Baltic or the major allele frequency changed by at least 0.1. As a quality filter, we required the SNP to be present in at least three individuals per population. To identify SNPs that show changes in allele frequency in both comparisons of the Atlantic to the Baltic Sea and the Danish Straits to the Baltic Sea, only SNPs identified by both comparisons were considered. This resulted in a list of SNPs per species that show changes in the Baltic Sea (Figure 4c).

To assess convergence between both species, we used the mapping of orthologous SNPs to identify which of the SNPs showing allele frequency changes in the Baltic were present in both species. Convergent SNPs were identified by filtering which of these SNPs show the same major allele in the Baltic Sea. Using the annotation of the SNPs provides information on their position inside genes and associated VEP consequences.

## Acknowledgements

We thank the Tvärminne Zoological Station (Laura Kauppi, Anna Vesanen) and Station Marine d’Arcachon (Xavier de Mautondouin) for collecting samples and the Kristineberg Marine Research Station (Leon Green, Elena Tamarit Castro, Linus Hammar Perry) and Ar Research Station on Gotland for help in sampling the pipefishes. This work was supported by a research grant (42153) from VILLUM FONDEN to JS, in addition to support by the University of Copenhagen Department of Biology’s “EcoCluster”.

DeiC (DeiC-KU-N2-2025160) to JD, and AQUASERV grant (PID: 37076) to JD. Lab work and genome assembly for the *Syngnathus typhle* reference genome was made possible through the Yggdrasil project funded by Carlsbergfondet Research Infrastructure Grant (CF22-0680) to MTPG. ET was supported by a Fulbright Scholarship.

## Data availability

All data for the *Syngnathus typhle* genome assembly are available on NCBI (PRJNA1226604) with accession numbers GCA_048301445.1 and GCA_048301605.1 for the two haplotypes. The genome is associated with BioProjects PRJNA1148218 for haplotype 1, which includes the whole genome nucleotide and protein sequences and PRJNA1148217 for haplotype 2, which includes the whole genome nucleotide sequences. The sample used for the genome has the BioSample ID SAMN36735486 (YG002_04). Illumina, Hi-C and HiFi sequencing data are available under PRJNA1393323, while PRJNA1393322 contains transcriptome sequencing data for gills and eyes. The VGP annotation for haplotype 1 is available under accession GCA_048301445.1-GB_2025_08_16.

Reads per sample for the population-level resequencing data is available on ENA under PRJEB107179 with individual ENA sample IDs given in the supplement (Table S2).

Assembly and annotation pipelines used for the *S. typhle* genome are available on GitHub (https://github.com/mahajrod/AssemblyBrute). All used scripts and data for the population-level and convergence analyses will be available upon publication.

## Notes

### Competing Interest Statement

The authors have declared no competing interest.

