## Supplementary Material for "Convergent molecular changes in populations of two pipefish species in the Baltic Sea"

**Table S1**: Sampling information summarized per location. Provided as a separate supplementary file in xlsx format (TableS1.xlsx).

**Table S2**: Sampling information, read counts as well as trimming and mapping statistics per individual. Provided as a separate supplementary file in xlsx format (TableS2.xlsx).

**Table S3**: SNP counts for the two species in different data sets and counts for SNP annotations in relation to genes. Abbreviations: *S. typhle*: *Syngnathus typhle*, *N. ophidion*: *Nerophis ophidion*, LD: linkage disequilibrium.

| **Data set** | **SNP count** |
|---|---:|
| SNPs total *S. typhle* | 1,488,252 |
| SNPs total *N. ophidion* | 19,565,453 |
| SNPs total *N. ophidion* (including chr 4) | 20,578,360 |
| SNPs not in LD *S. typhle* | 154,292 |
| SNPs not in LD *N. ophidion* | 1,504,488 |
| SNPs 1 per 10kb window *S. typhle* | 31,795 |
| SNPs 1 per 10kb window *N. ophidion* | 166,496 |
| SNPs in genes *S. typhle* | 1,146,984 |
| SNPs in genes *N. ophidion* | 12,723,245 |
| SNPs in exons *S. typhle* | 264,222 |
| SNPs in exons *N. ophidion* | 637,432 |
| SNPs convergent *S. typhle* and *N. ophidion* | 4,168 |
| Genes convergent *S. typhle* and *N. ophidion* | 17,513 |

**Table S4**: List of convergent genes including detailed information on the gene's annotation and location in the genome for each of the two species *Syngnathus typhle* and *Nerophis ophidion*. Provided as a separate supplementary file in xlsx format (TableS4.xlsx).

**Table S5**: Five convergent SNPs including their annotation and statistics on $F_{ST}$, $d_{XY}$ and $\pi$ for each species and each population comparison and allele frequencies for each population and species. Provided as a separate supplementary file in xlsx format (TableS5.xlsx).

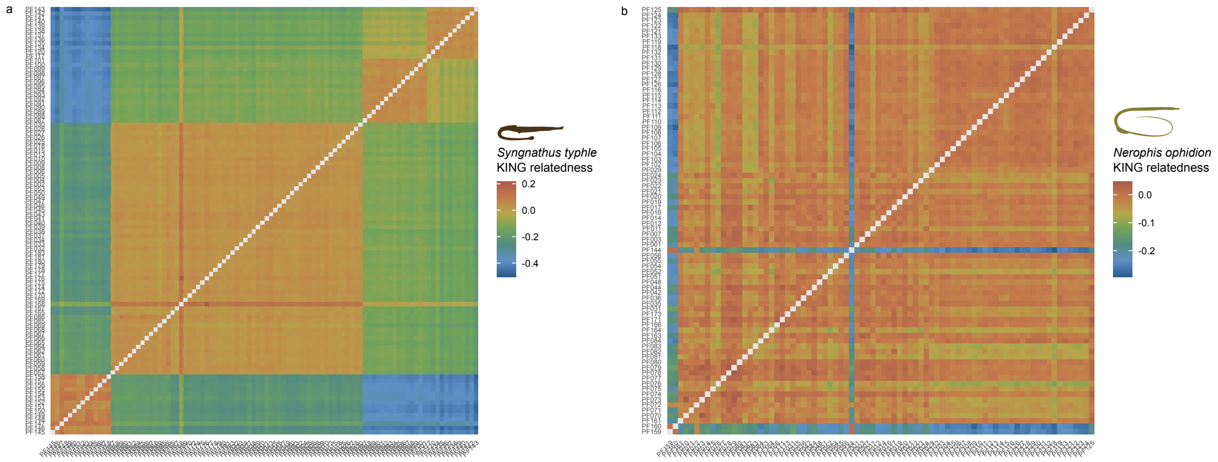


**Figure S1**: Reasoning for the removal of one individual (PF168) of the broadnosed pipefish (*Syngnathus typhle*) from Guldborgsund. (a) KING relatedness score for each combination of individuals of *S. typhle*. One individual clearly stands out with high relatedness to all other individuals. (b) Same plot as (a) but for the straightnose pipefish (*Nerophis ophidion*). Overall relatedness is lower and no outliers with elevated KING score can be seen.

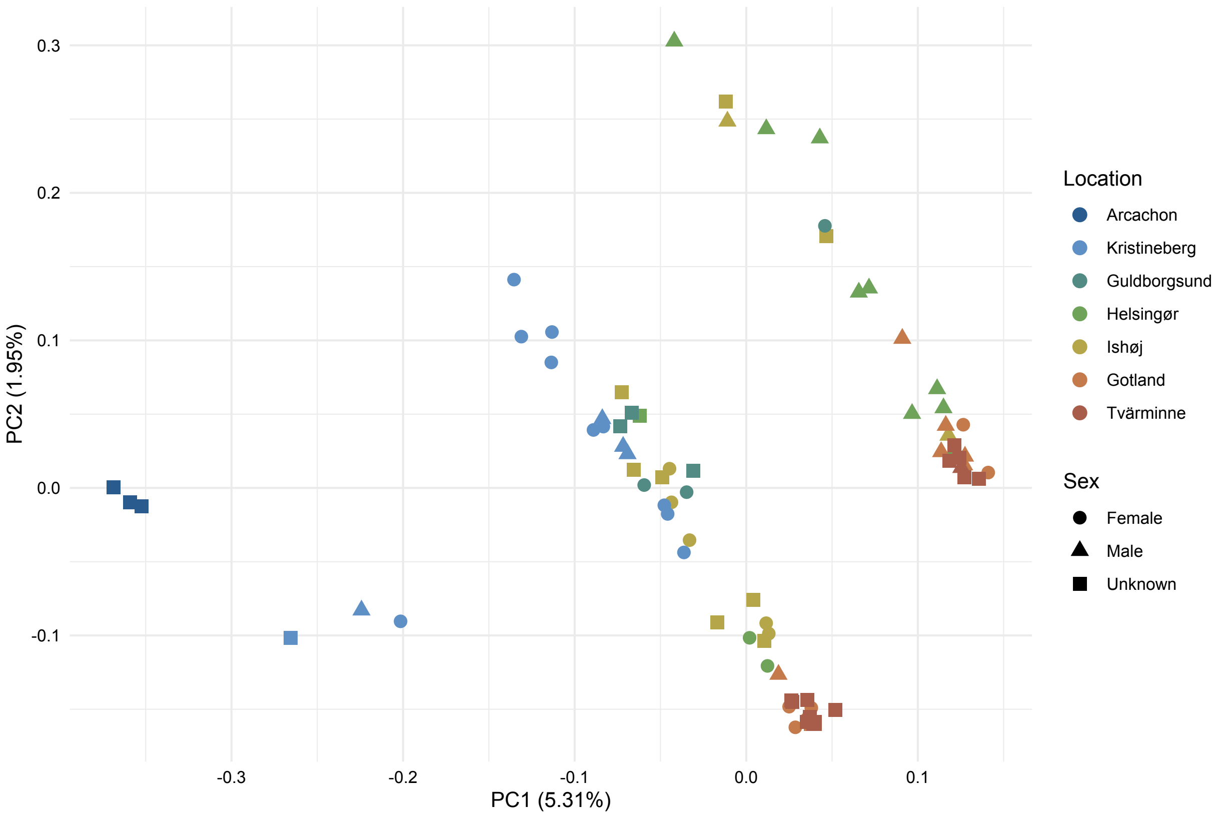


**Figure S2**: Reasoning for the removal of chromosome 4 (NC_084614.1) of the straightnose pipefish (*Nerophis ophidion*). (a) PCA plot of all SNPs in *N. ophidion* including the SNPs on chromosome 4. Three lines can be seen that do not correspond to locations indicated by color but seem to match better with the sex indicated by the shape (b) Linkage map for chromosome 4, which shows high linkage disequilibrium (LD) across the whole chromosome. (c) Linkage map for chromosome 1 as a representative of all other chromosomes, which in comparison to chromosome 4 shows less widespread LD.

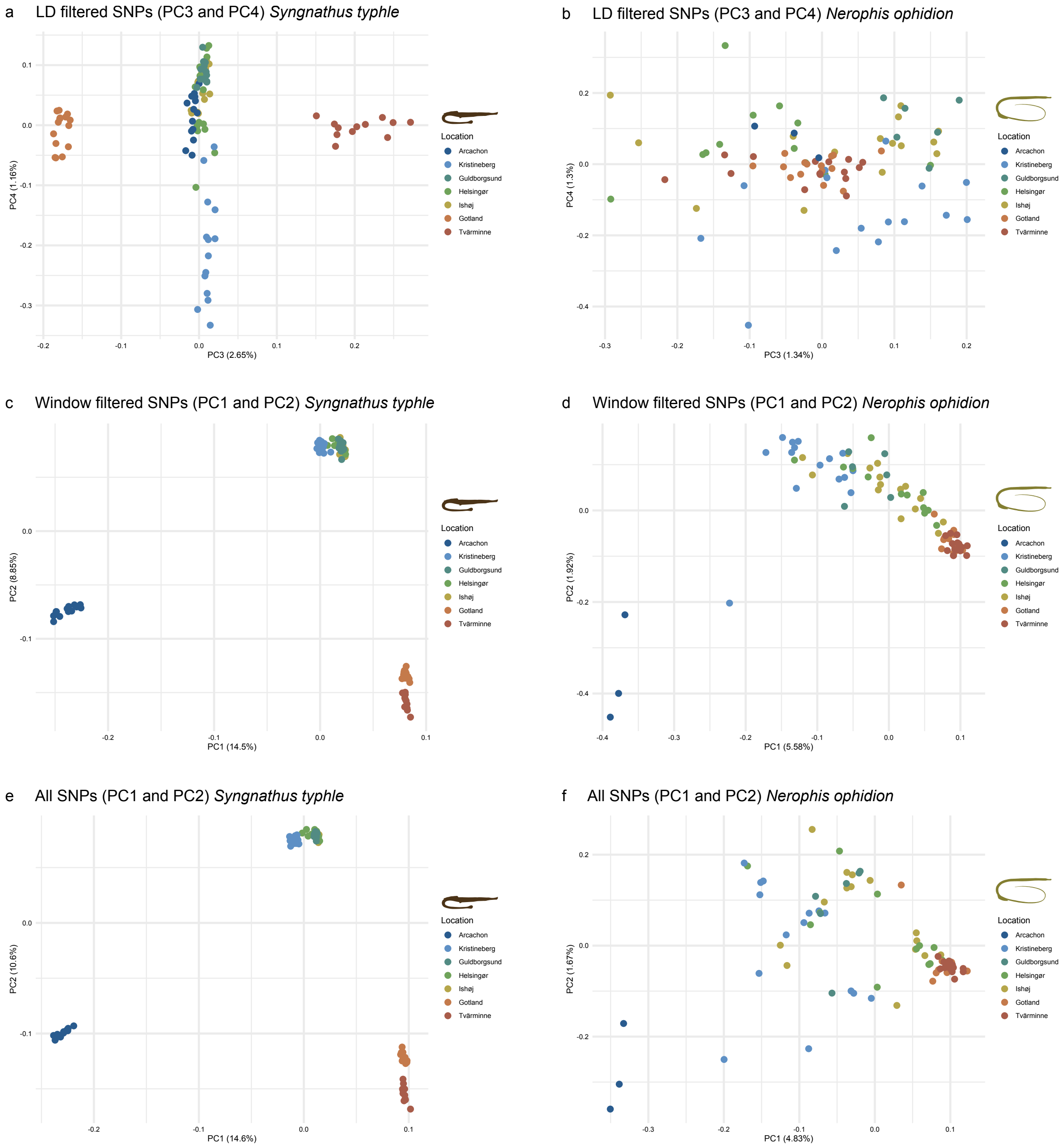


**Figure S3**: Additional PCA plots for different data sets. Plots for the broadnosed pipefish (*Syngnathus typhle*) are shown on the left, plots for the straightnose pipefish (*Nerophis ophidion*) on the right. (a, b): PC3 against PC4 for data filtered for linkage disequilibrium (LD). The data set corresponds to the data used in Figure 2a and Figure 2d. (c, d): Window LD filtered data set, where one random SNP is kept per 10 kb window. Patterns are largely similar to Figure 2a and Figure 2b. (e, f): PCA using all SNPs, without an LD filter. Patterns for *S. typhle* are largely similar, but PCs have more weight. Patterns in *N. ophidion* are not as clear, as there are a very large number of linked SNPs that outweigh the others.

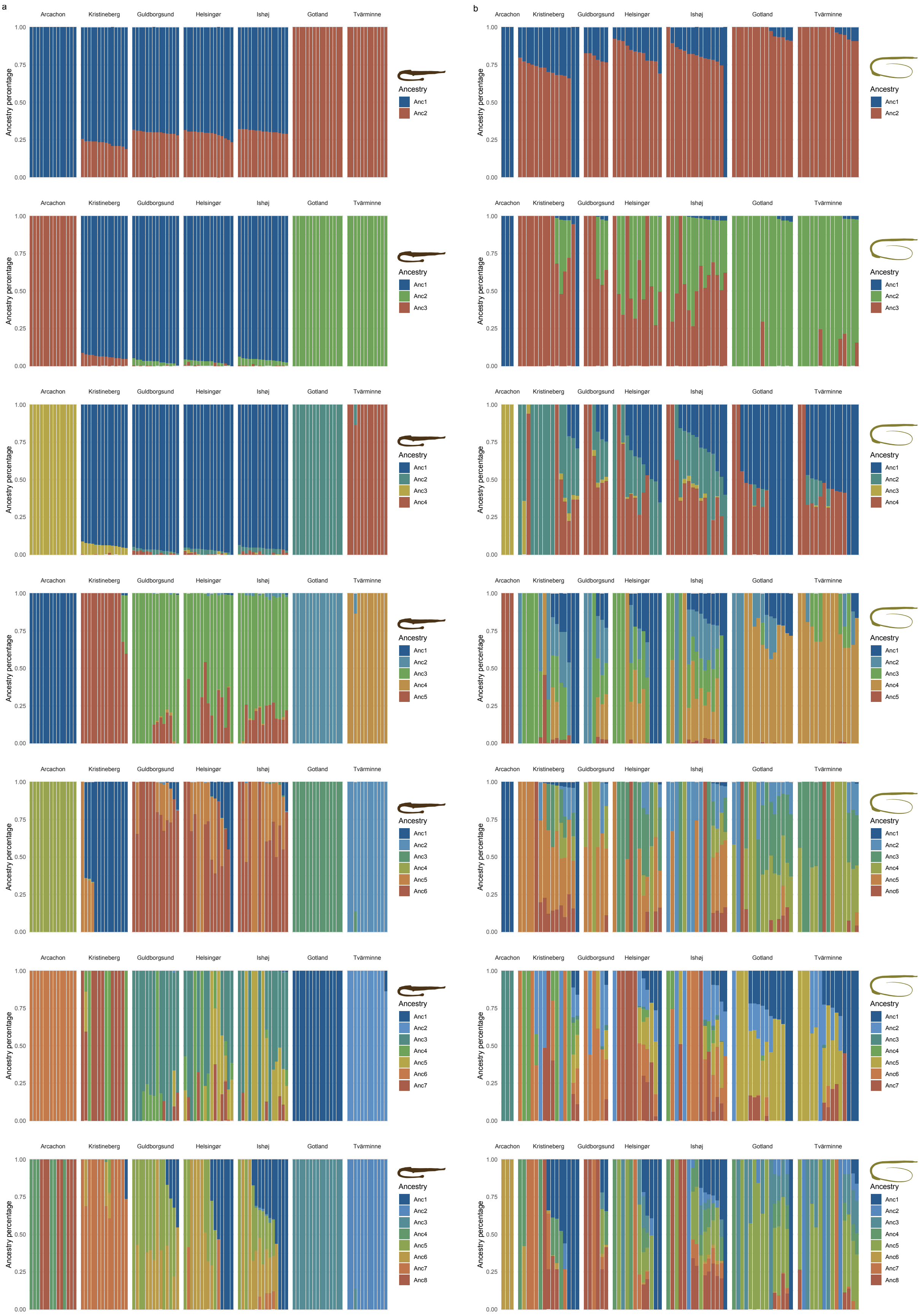


**Figure S4**: Admixture plots for k ranging from 2 to 8 for (a) the broadnosed pipefish (*Syngnathus typhle*) and (b) the straightnose pipefish (*Nerophis ophidion*) on the right.

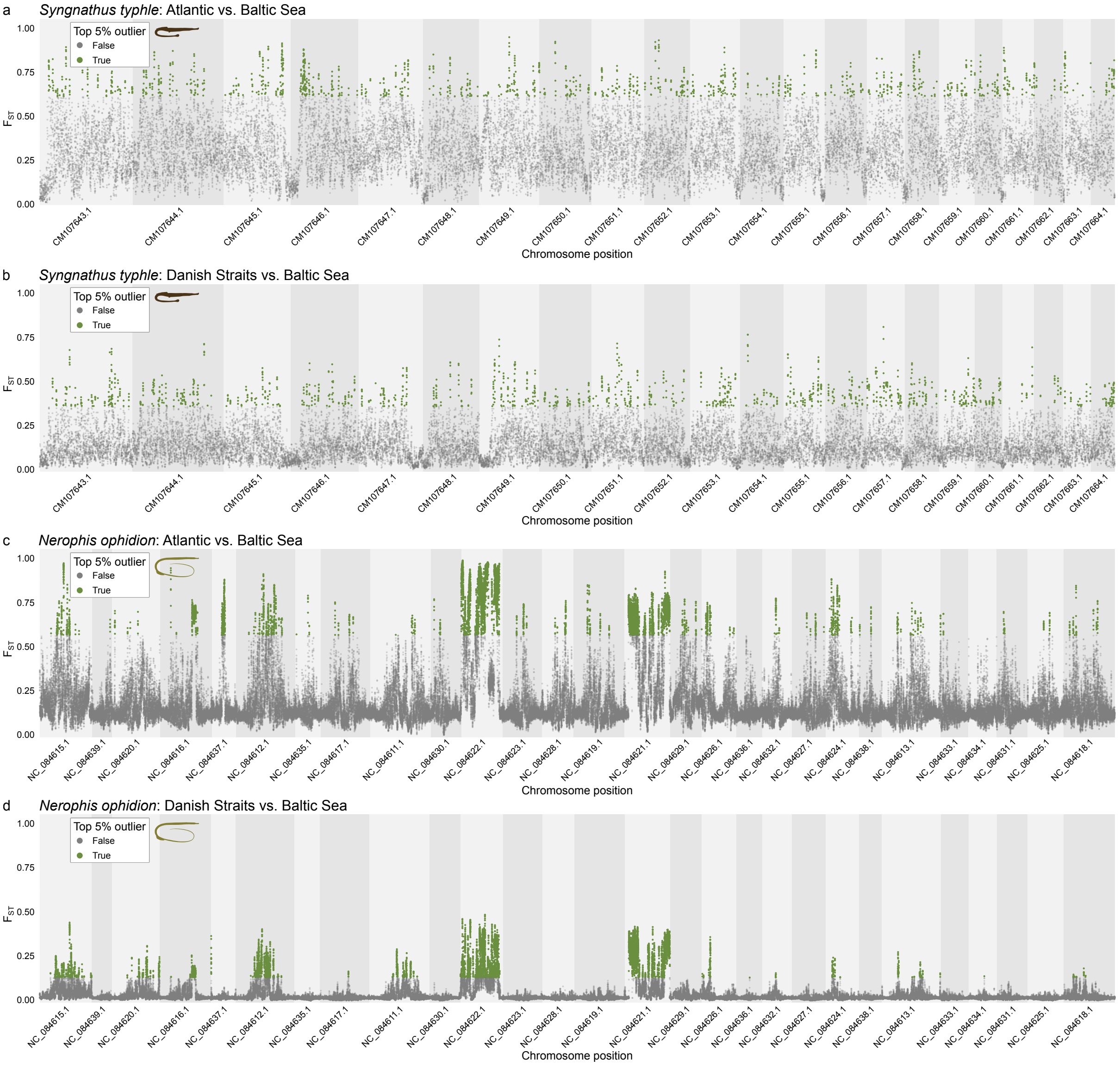


**Figure S5**: Chromosome-level differentiation of the broadnosed pipefish (*Syngnathus typhle*) and the straightnose pipefish (*Nerophis ophidion*) using $F_{ST}$ windows across the whole genome. (a) Manhattan plot showing $F_{ST}$ values across the whole genome between Atlantic and Baltic Sea populations of *S. typhle*. The top 5% outliers are colored in green. (b) Same plot as (a) but for Danish Straits and Baltic populations. (c) Same plot as (a) but for *N. ophidion*. (d) Same plot as (b) but for *N. ophidion*. Chromosomes are ordered according to their syntenic relationships with *S. typhle* (synteny of all chromosomes shown in Figure S7).

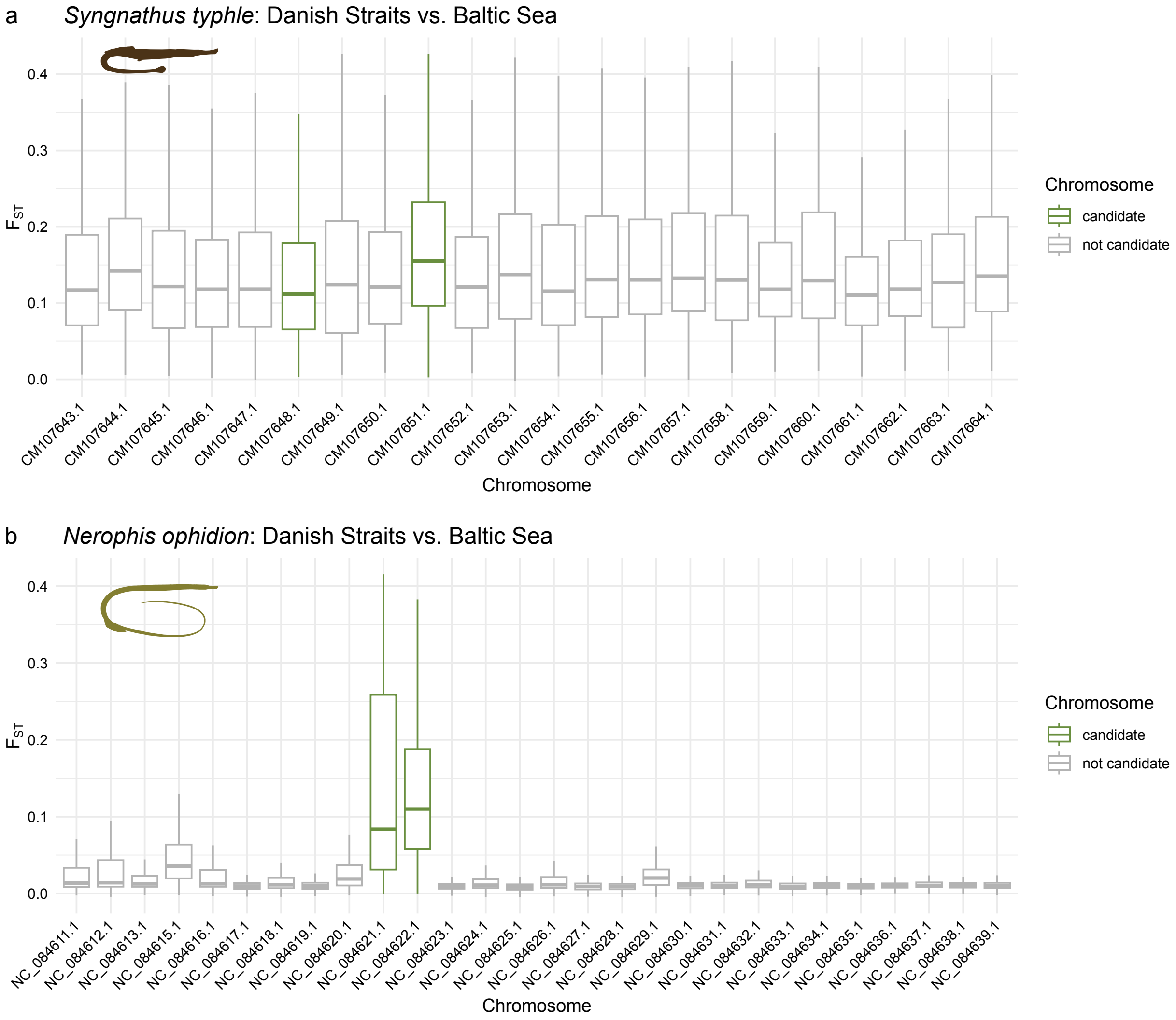


**Figure S6**: Average $F_{ST}$ pattern per chromosome between the Danish Straits and the Baltic Sea in (a) the broadnosed pipefish (*Syngnathus typhle*) and (b) the straightnose pipefish (*Nerophis ophidion*).

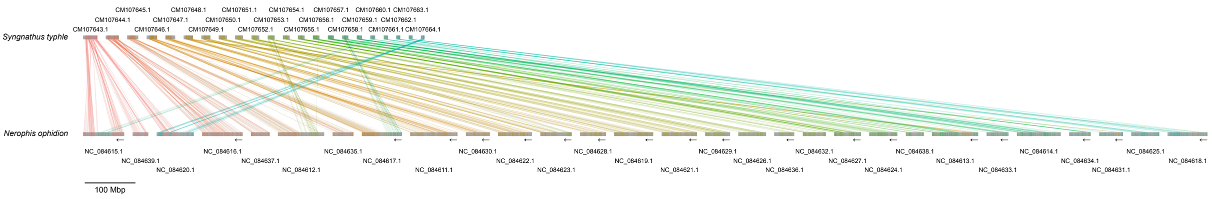


**Figure S7**: Synteny of chromosomes all between the broadnosed pipefish (*Syngnathus typhle*) and the straightnose pipefish (*Nerophis ophidion*). Reverse complemented sequences in *N. ophidion* are indicated with arrows.

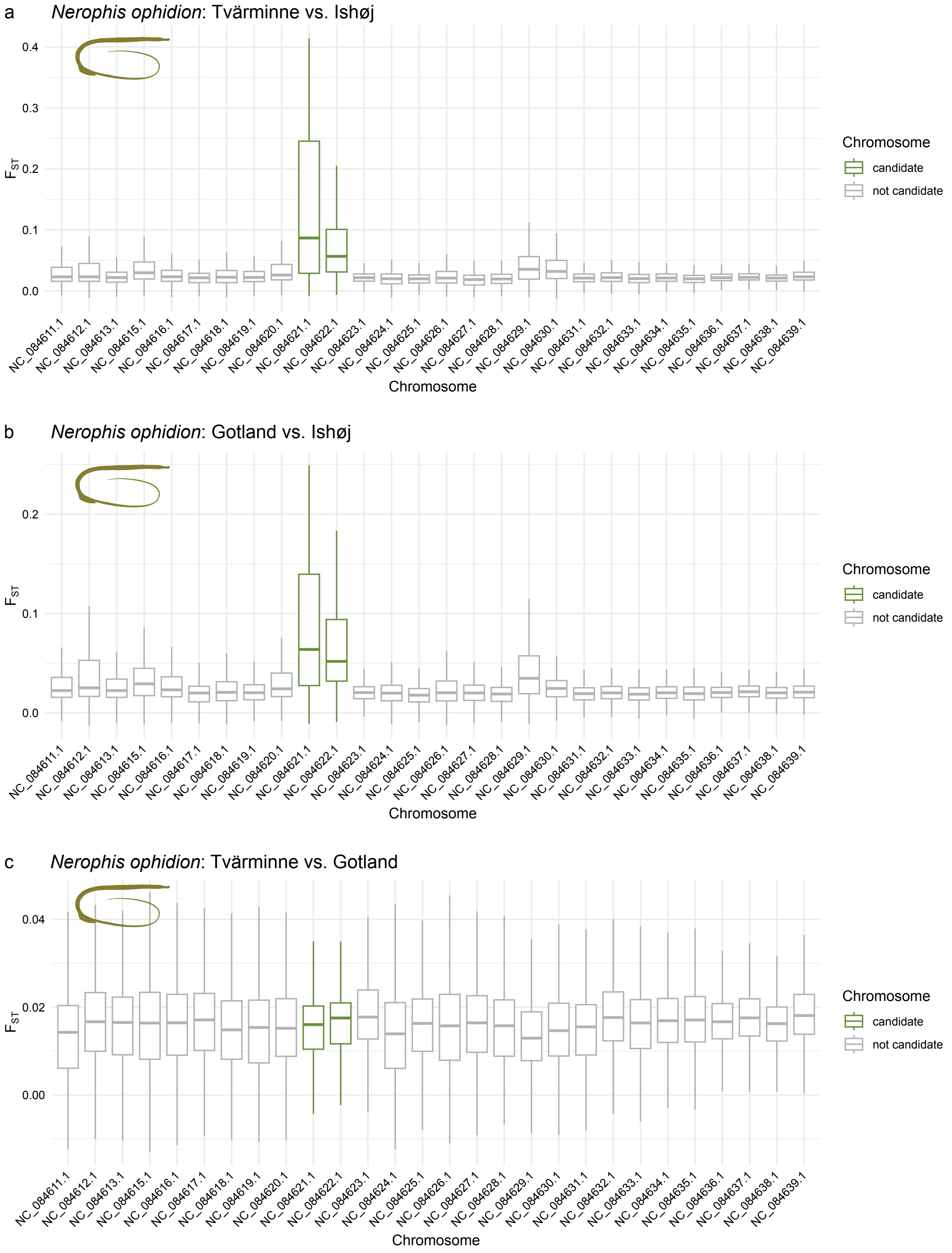


**Figure S8**: Shared elevation of $F_{ST}$ in the straightnose pipefish (*Nerophis ophidion*) in both populations of the Baltic Sea. (a) Comparison of Tvärminne (Baltic) to Ishøj (Danish Straits) shows higher $F_{ST}$ in chromosomes NC_084621.1 and NC_084622.1 compared to all other chromosomes. (b) Same plot as a) but comparing Gotland (Baltic) to Ishøj (Danish Straits). c) Comparison of Tvärminne (Baltic) to Gotland (Baltic) does not show higher $F_{ST}$ in chromosomes NC_084621.1 and NC_084622.1 compared to all other chromosomes.

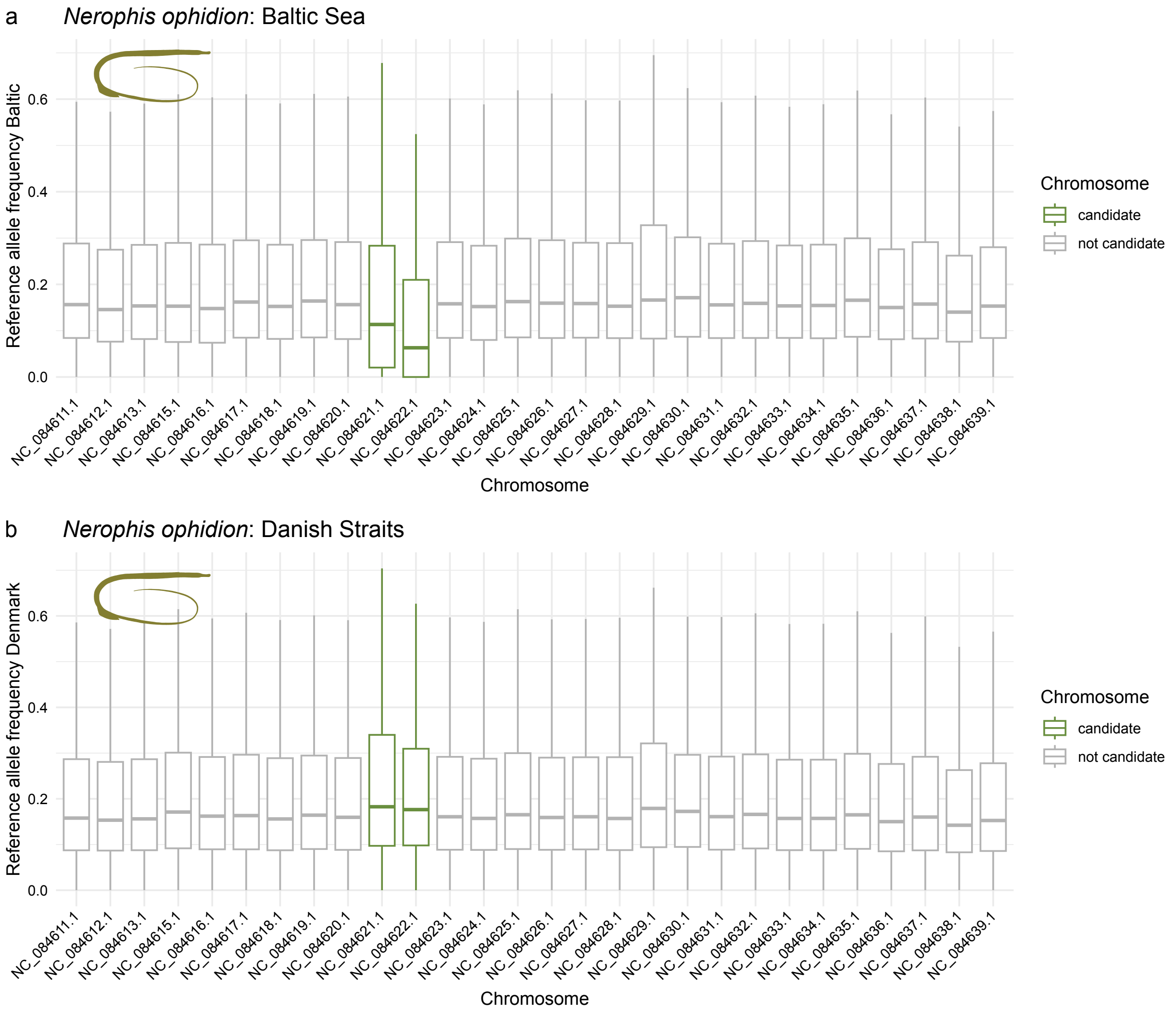


**Figure S9**: Allele frequency pattern across chromosomes in the straightnose pipefish (*Nerophis ophidion*). (a) Boxplot of the allele frequency of reference allele per chromosome in the Baltic populations, (b) Boxplot of the allele frequency of reference allele per chromosome in the Danish Straits populations.

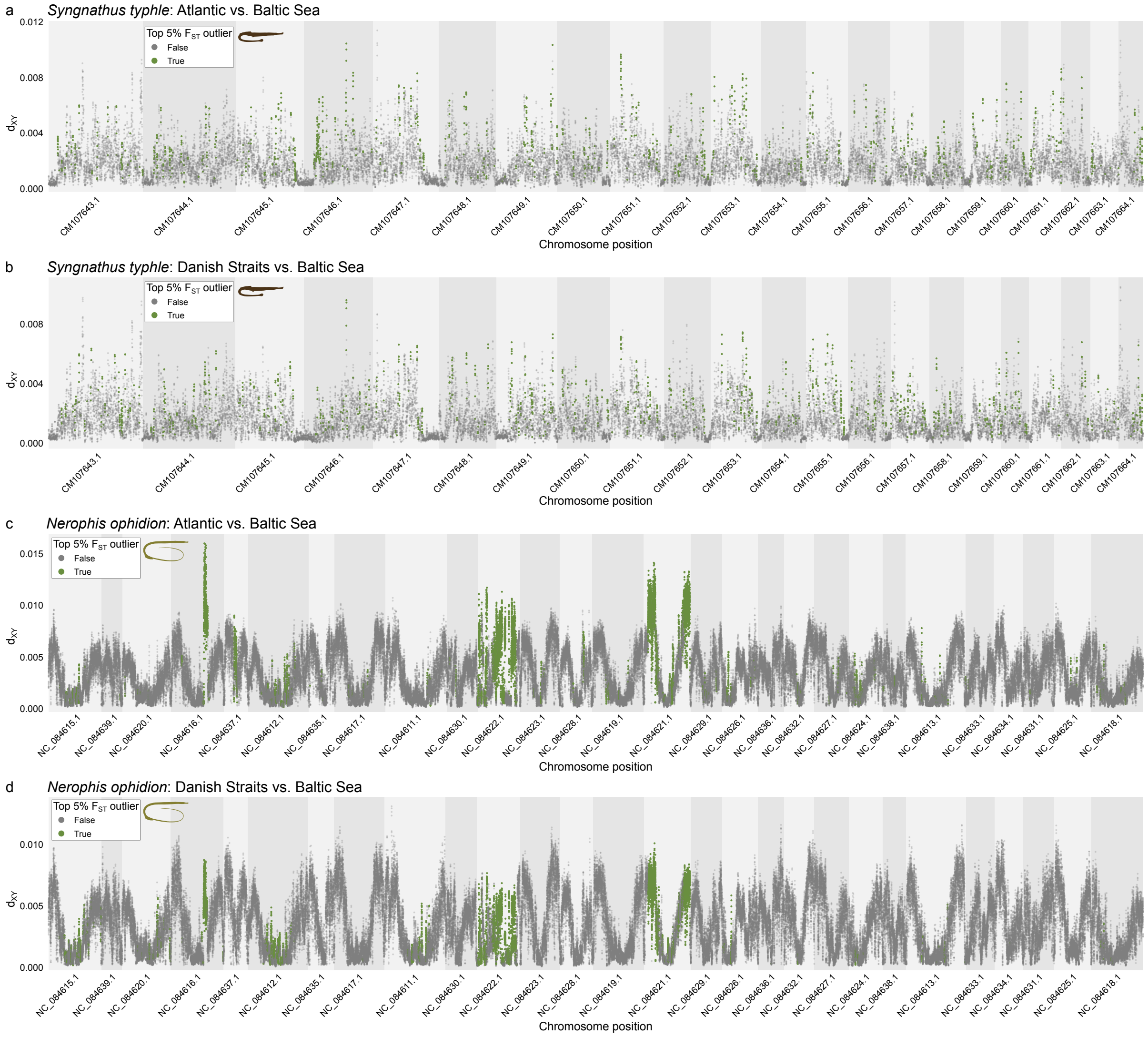


**Figure S10**: Chromosome-level divergence of the broadnosed pipefish (*Syngnathus typhle*) and the straightnose pipefish (*Nerophis ophidion*) using absolute genetic divergence ($d_{XY}$) windows across the whole genome. (a) Manhattan plot showing $d_{XY}$ values across the whole genome between Atlantic and Baltic Sea populations of *S. typhle*. Colors are based on $F_{ST}$ values for each window, with the top 5% outliers colored in green. (b) Same plot as (a) but for Danish Straits and Baltic populations. (c) Same plot as (a) but for *N. ophidion*. (d) Same plot as (b) but for *N. ophidion*. Chromosomes are ordered according to their syntenic relationships with *S. typhle* (synteny of all chromosomes shown in Figure S7).

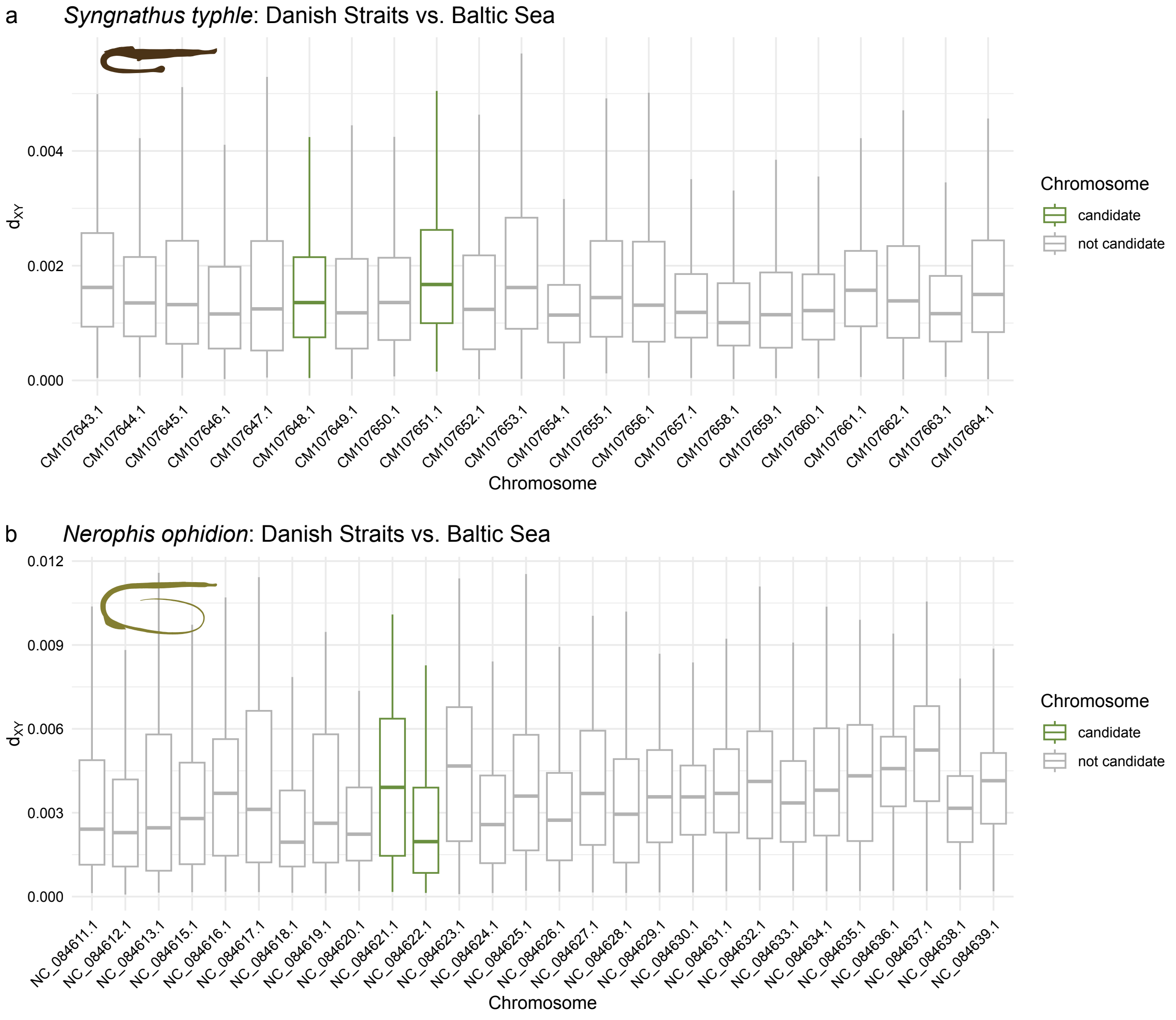


**Figure S11**: Average absolute genetic divergence (d$_{XY}$) across chromosomes between the Danish Straits and the Baltic Sea in (a) the broadnosed pipefish (*Syngnathus typhle*) and (b) the straightnose pipefish (*Nerophis ophidion*).

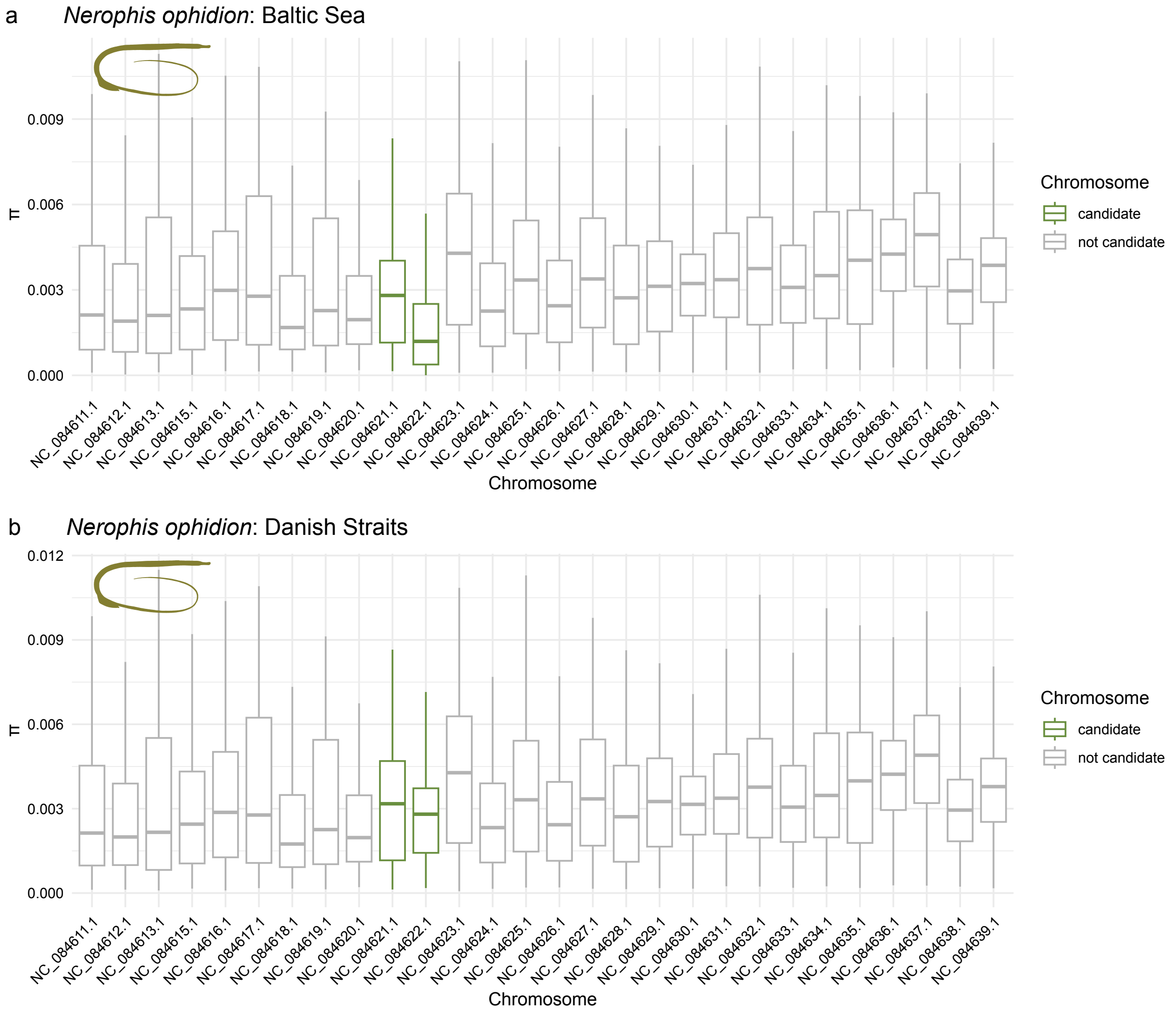


**Figure S12**: Nucleotide diversity (π) across chromosomes in the straightnose pipefish (*Nerophis ophidion*). (a) Boxplot of the allele frequency of reference allele per chromosome in the Baltic populations. (b) Boxplot of the allele frequency of reference allele per chromosome in the Danish Straits populations.

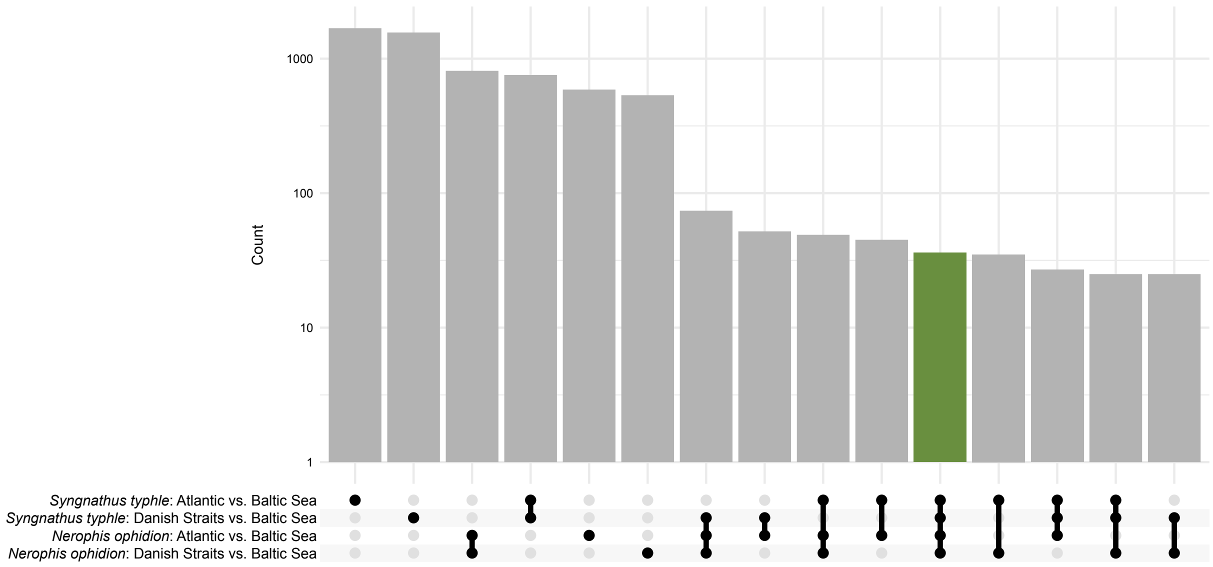


**Figure S13**: Upset plot that shows how many genes overlap when comparing the four different comparisons (Atlantic-Baltic Sea and Danish Straits-Baltic Sea for each species, the broadnosed pipefish (*Syngnathus typhle*) and the straightnose pipefish (*Nerophis ophidion*)) based on $F_{ST}$ outliers. Genes identified in all comparisons are highlighted in green.

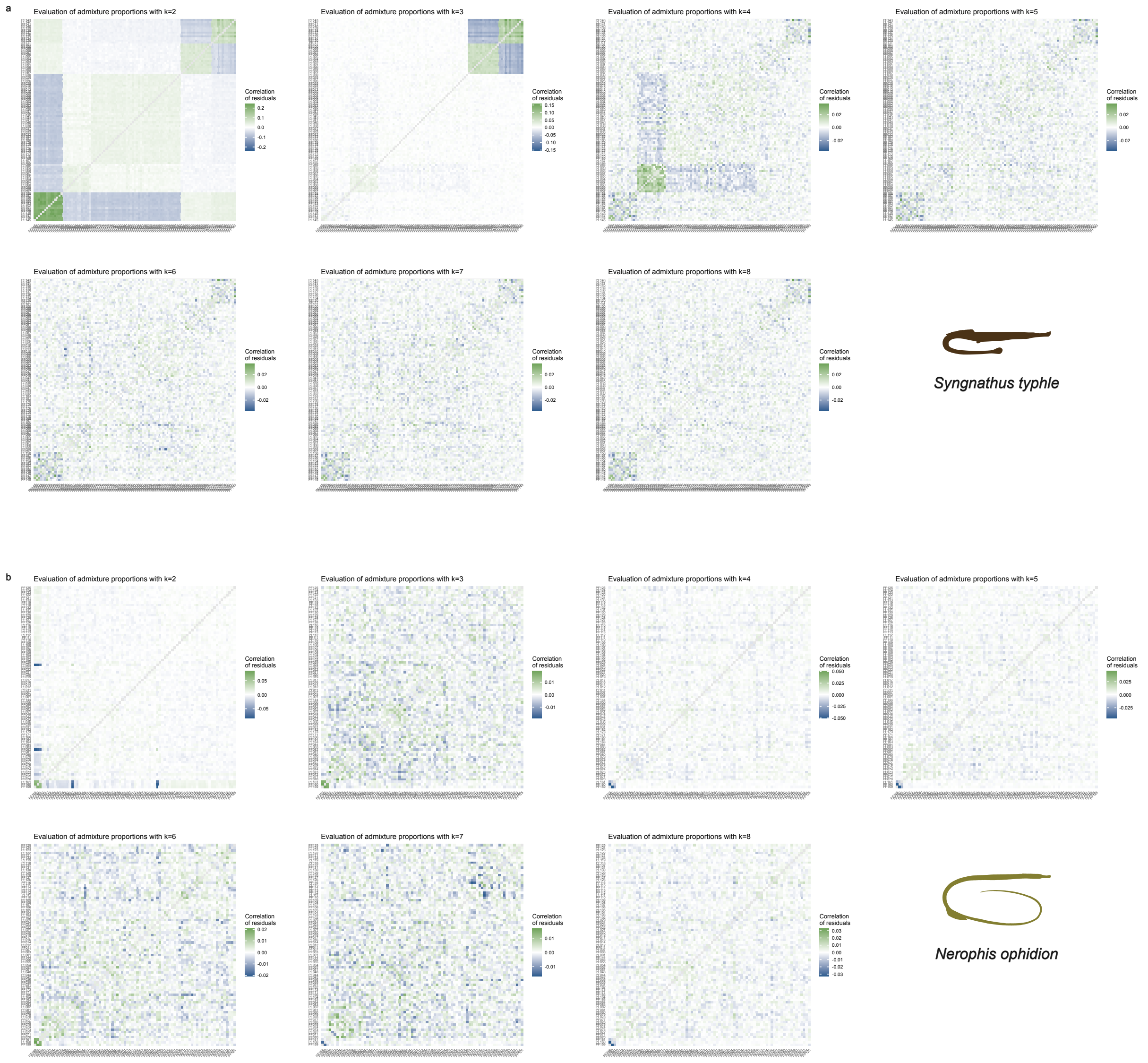


**Figure S14**: Admixture residuals for (a) the broadnosed pipefish (*Syngnathus typhle*) and (b) the straightnose pipefish (*Nerophis ophidion*) for major mode of NGSadmix runs for k=2-8.
